# MYO1F interactome reveals the SH3-domain linked CASS complex at podosomes and the phagocytic cup

**DOI:** 10.1101/2025.07.04.663186

**Authors:** Susan D. Arden, Eva Pennink, András Lakatos, Gillian M. Griffiths, Anna H. Lippert, Folma Buss

## Abstract

MYO1F, a long-tailed myosin of class I, is selectively expressed in immune cells and upregulated in microglia associated with neurodegenerative pathogenesis. The intracellular functions of myosin motors involve adaptor proteins, which regulate cargo attachment and intracellular motor recruitment. To define the MYO1F interactome, we performed an in situ proximity labelling-based proteomic analysis in human myeloid U937 cells. We identified a distinct SH3-domain-dependent adaptor module comprising <u>C</u>D2AP, <u>A</u>SAP1, <u>S</u>H3BP2, and <u>S</u>H3KBP1 (CASS complex), which localizes with MYO1F in podosomes and phagocytic cups. Structural modelling and mutagenesis confirmed multivalent proline-rich motif interactions of the CASS complex with the MYO1F SH3 domain. Further deletions revealed a second group of membrane-associated adaptor proteins that bind to the MYO1F pleckstrin homology (PH) domain. Immunofluorescence in macrophages and microglia confirmed the conserved localization of MYO1F and its adaptors at actin-rich podosomes and phagocytic cups. Functional assays demonstrated that MYO1F recruitment to the phagocytic cup requires motor activity and intact PH and SH3 domains. This study provides the first comparative interactome of MYO1F and its paralogue MYO1E and supports a role for MYO1F in podosomes and during phagocytosis in both peripheral and brain-resident myeloid cells.

## INTRODUCTION

Within eukaryotic cells, myosin motor proteins move cargo over short distances along actin tracks, regulate plasma membrane dynamics through the actin cortex and provide flexible tethering of organelles, vesicles and protein complexes to the actin^1,2,3^. Myosins of class I are widely expressed monomeric single headed motors^4^. The biological functions of class I myosins are at the membrane-cytoskeleton interface, providing mechanical force and tension that allows remodelling of membranes relative to the underlying actin network. They play important roles in plasma membrane dynamics and membrane trafficking as well as in the organisation of the actin cytoskeleton ^5,6^.

In humans a total of 8 class I myosins are expressed (MYO1A-MYO1H), which are subdivided into the short-tailed and the long-tailed myosins ^7^. All myosins of class I can directly interact with cell membranes through the pleckstrin homology domain (PH) present in the tail homology 1 (TH1) domain ^8^. Two additional domains are present only in the long-tailed MYO1E and F, a proline-rich tail homology 2 (TH2) and a C-terminal SH3-domain (SH3) ^9,10^. MYO1E can be found in a wide variety of cell types and tissues with enriched expression in the kidney and intestine^11,9^ (www.proteinatlas.com), while MYO1F expression is restricted to blood and immune cells including peripheral lymphoid and myeloid cells but also brain-resident microglia^12, 13, 14, 15^.

Microglia are the primary immune cells of the central nervous system, playing a crucial role in the pathophysiology of various neurodegenerative diseases ^16^. In both human Alzheimer’s disease (AD) patients as well as AD and frontotemporal dementia/amyotrophic lateral sclerosis (FTD/ALS) murine models, MYO1F is significantly upregulated in microglia compared to healthy controls. This alteration in MYO1F expression parallels changes observed in established AD and FTD/ALS-associated risk genes, such as TREM2 and GRN^15,17^. Furthermore, whole-genome gene-expression profiling has identified MYO1F as a key node within immune-related networks in patients with late-onset AD^14^. Additionally, systems biology analysis suggests that MYO1F is one of the seven microglia-specific genes potentially involved in a shared molecular pathway underlying the human ageing process and multiple neurodegenerative diseases such as AD, Parkinson’s disease, and Huntington’s disease^18^.

In peripheral myeloid cells MYO1F deficiency results in increased adhesion and reduced motility of neutrophils in vitro^12^ and a significant reduction in extravasation, attributed to the neutrophils’ incapacity to deform their nucleus while squeezing through endothelial cell barriers^13^. Another study shows that MYO1F mediates intercellular adhesion, which allows intestinal macrophage differentiation to a M1-phenotype. Importantly, in a mouse model of colitis, the absence of MYO1F causes a reduction in proinflammatory cytokine secretion, less tissue damage and increased tissue repair ^19^.

MYO1E and MYO1F are present in podosomes ^20^ and invadopodia ^21^, which are micron-sized, actin-rich structures critical for cell adhesion, matrix degradation, and mechanosensing ^22, 23^. While invadosomes are a feature of invasive cells ^24^, podosomes are predominantly found in monocytic cells ^25^ but are also observed in smooth muscle ^26^ and endothelial cells ^27^. MYO1E has been shown to negatively regulate actin polymerization at the ventral layer of podosomes ^28^. Interestingly, actin-rich, podosome-like structures have also been identified at the leading edge of the phagocytic cup ^29, 30^. These phagocytic podosomes, which contain both MYO1E and MYO1F, are thought to tether the plasma membrane to the underlying actin cortex, thereby increasing membrane tension ^31,32^. The absence of MYO1F and/or MYO1E results in increased actin polymerization, leading to densely packed actin filaments in the phagocytic cup which subsequently slow closure of the phagocytic cup ^33^.

In this study, we employ in situ proximity labelling using BioID to characterize the overall interactome of both MYO1F and MYO1E. Our analysis identifies binding partners common to both myosins across different cell types as well as a number of protein complexes unique to either MYO1F or MYO1E. Comparative analysis of these in vivo proximity maps highlights a functional overlap between these two myosins. We subsequently focus on MYO1F and conduct affinity pull-down assays and co-localization studies to validate the MYO1F binding partners identified in BioID experiments. Using a series of MYO1F deletion mutants lacking either the SH3 domain (ΔSH3) or with point mutations in the PH domain (ΔPH) ^34^, which abolishes membrane binding, we determined the essential binding domains for different adaptor proteins. Our functional studies uncovered novel binding partners, such as ArfGAP With SH3 Domain, Ankyrin Repeat And PH Domain 1 (ASAP1)^35,36^, SH3 Domain Containing Kinase Binding Protein 1 (SH3KBP1)^37,38^, CD2 Associated Protein (CD2AP)^38,39^ as well as the known binding partner SH3 Domain Binding Protein 2 (SH3BP2)^40,41,42^, which either colocalise with MYO1F in ventral podosomes in macrophages and microglia, or in the phagocytic cup during phagocytosis. This work underscores the diverse interaction network of MYO1F, providing new insights into the molecular mechanisms that enable this multifunctional motor protein to participate in a number of cellular processes. Notably, our findings also give insight into potential roles for MYO1F in neurodegeneration, as mutations in two of its adaptor proteins, CD2AP and SH3KBP1, have been implicated in late-onset AD.

## RESULTS

### Identification of the MYO1F interactome using in situ proximity labelling

To shed further light on the various proposed functions of MYO1F in myeloid cells and to capture potential MYO1F binding partners involved in these processes, we used in situ proximity labelling in living cells under steady state conditions. MYO1F is specifically expressed in myeloid cells such as THP-1, U937 and microglia while MYO1E is expressed in a wider variety of cell types and tissues including retinal epithelial cells (RPE) (Figure 1A). For our *in situ* proximity labelling experiments, we generated U937 cell lines stably expressing the MYO1F tail domain (aa 692-1098), which includes the PH-domain in TH1 and the C-terminal SH3-domain, tagged at the N-terminus with the promiscuous biotin ligase, BirA R118G (BirA*) (Figure 1B). We verified expression of the BirA*-MYO1F tail by immunoblotting (Supplementary Figure 1A). After labelling with biotin for 24 hours, biotinylated proteins were collected by streptavidin pull-down and enriched proteins identified by mass spectrometry. Results from 6 independent experiments for BirA*-MYO1F tail and 3 each for both mutants, BirA*-MYO1F tail ΔPH or BirA*-MYO1F tail ΔSH3 were analysed against 6 BirA*- only U937 control pulldowns using label-free quantification (LFQ) intensities (Figure 1). Using a difference ≥ 4 of MYO1F over control we identified 47 significantly enriched proximal proteins shown in figure 1C as a heat map of the relative abundance of each protein interaction with MYO1F. These include the only known binding partner of MYO1F, SH3BP2 ^42^, and more than 40 novel MYO1F-associated proteins which might bind either directly or as part of larger MYO1F-asociated protein complexes (Figure 1C and 1D).

**Figure 1:**
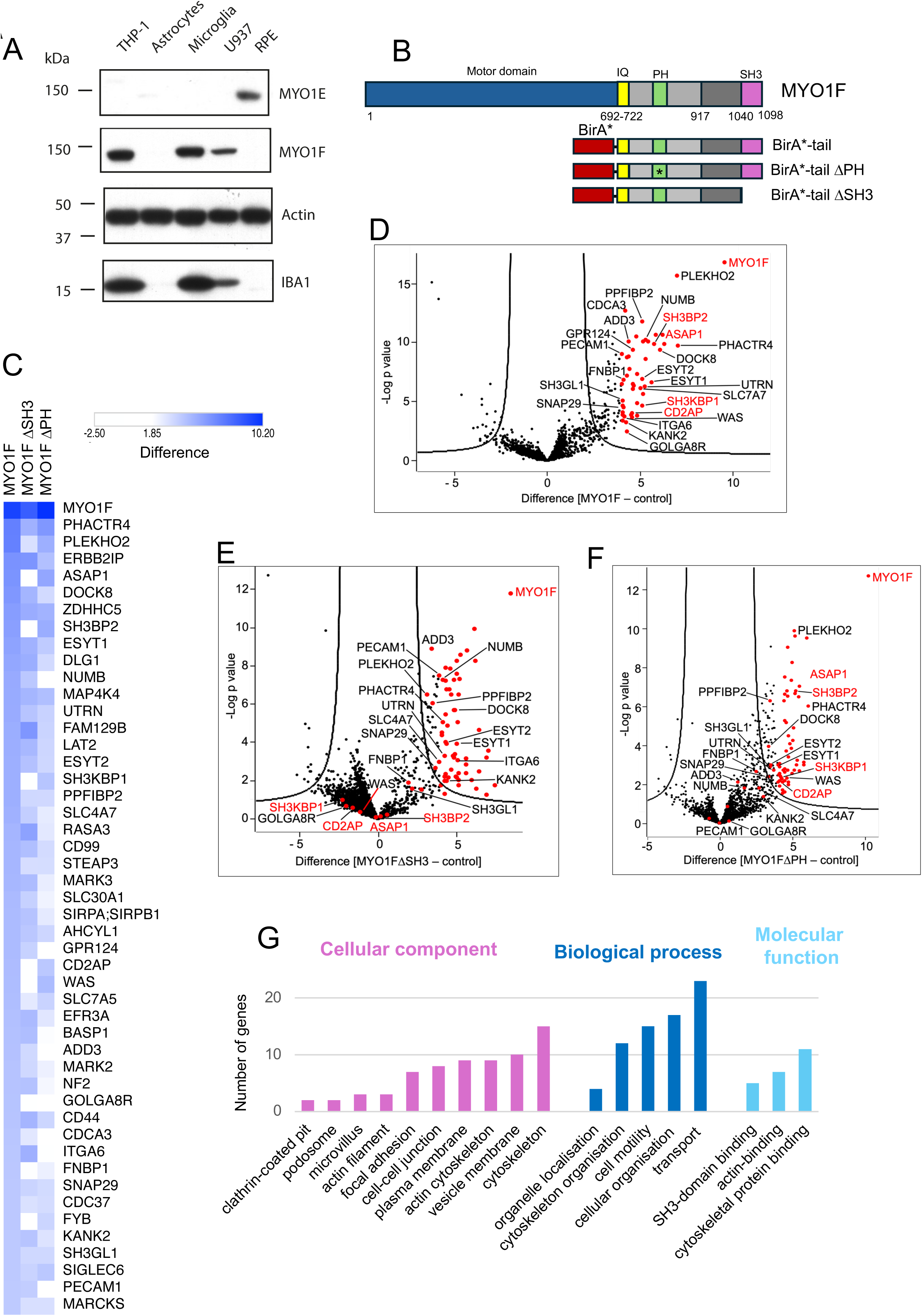
In situ proximity labelling reveals new binding partners for MYO1F. **A**. Expression of MYO1E and MYO1F in different cell types shown by Western blotting with actin as a loading control. Iba-1 was used to identify cells of monocytic lineage such as THP-1, U937 and microglia. **B**. Schematic diagram of BirA*-MYO1F tail domain and mutant constructs ΔPH or ΔSH3 domain. **C**. Heat map of abundance of protein interactions with MYO1F, MYO1FΔSH3 or MYO1FΔPH determined by BioID was generated using Morpheus software (https://software.broadinstitute.org/ morpheus). Shown are protein interactions with a difference ≥ 4 over control. Data is shown from 6 independent experiments for MYO1F, and 3 independent experiments for the two mutants. **D**. Volcano plot of LFQ data from BirA*-MYO1F-tail over BirA* control from U937-stable cell lines. The negative logarithmic *P* value was plotted against the t-test difference (n=6). The hyperbolic cutoff curve delimitates significantly enriched proteins from common hits (FDR of 0.01 and SO of 2). Significantly enriched (red dots) and other notable proteins are labelled by name. Proteins of specific interest are labelled in red. The complete list is given in supplementary data. **E.** Volcano plot of LFQ data from BirA*-MYO1F-tailΔSH3 over BirA* control. The negative logarithmic *P* value was plotted against the t-test difference (n=3). The hyperbolic cutoff curve delimitates significantly enriched proteins from common hits (FDR of 0.01 and SO of 2). **F.** Volcano plot of LFQ data from BirA*-MYO1F-tail ΔPH over BirA* control. The negative logarithmic *P* value was plotted against the t-test difference (n=3). The hyperbolic cutoff curve delimitates significantly enriched proteins from common hits (FDR of 0.01 and SO of 2). **G**. Gene ontology (GO) classification of MYO1F interactome identified by in situ proximity labelling. GO cellular component, biological process and molecular function enrichment analyses were performed using ShinyGO software (Ge SX, Jung D and Yao R. Bioinformatics 36: 2628-2629, 2020). A term with P < 0.05 was considered to be significantly overrepresented.

To dissect the domain-specific interaction profiles of MYO1F, we compared the interactomes of the full-length MYO1F tail with its SH3-and PH-domain mutants (Figure 1C-F). This analysis revealed a core group of adaptor proteins whose interaction with MYO1F was dependent on the SH3 domain (Figure 1C and E). These include SH3BP2 (also called 3BP2), SH3KBP1(also called CIN85), CD2AP (also called CMS), and ASAP1. These proteins, characterized by proline-rich regions or SH3 domains themselves, form a distinct protein module we termed the CASS complex (CD2AP-ASAP1-SH3BP2-SH3KBP1). The MYO1F interactome contains three further proteins that bind to the SH3 domain, the Wiskott-Aldrich Syndrome protein (WASP)^43,44^, Formin-binding protein 1 (FNBP1)^45,46^, and FYN-binding protein (FYB)^47,48^ (Figure 1C and E). These proteins share common pathways and functions related to the actin cytoskeleton and immune cell signalling, and FYB and WASP have been identified as part of a molecular complex during phagocytosis in macrophages^48^. Interestingly, mutation of the PH domain in the TH1 region did not affect binding of the CASS complex, but reduced binding to proteins with membrane-associating functions, such as for example utrophin (UTRN)^49,50^ a cytoskeletal linker protein that anchors the actin cytoskeleton to the plasma membrane, ADD3 (Adducin gamma)^51^ a cytoskeletal protein that promotes spectrin-actin assembly at the plasma membrane, ITGA6 (Integrin alpha-6)^52^ and PECAM1^53,54^ an endothelial cell adhesion molecule (Figure 1C and F). These interactions likely depend on proper membrane localization of MYO1F through its PH domain and may represent a second class of functional proteins involved in plasma membrane anchoring and tension regulation.

To better understand the biological processes, molecular functions, and cellular localizations associated with the MYO1F interactome, we performed a Gene Ontology (GO) analysis on the 47 significantly enriched binding partners that were identified in our in situ labelling experiment (Figure 1G). The results show a significant enrichment for GO terms associated with cytoskeletal organization and membrane-cytoskeleton coupling and cellular component terms including vesicle membranes, focal adhesions and podosomes, reinforcing the proposed role of MYO1F at podosomes and during phagocytosis.

### Identification of the MYO1E interactome using in situ proximity labelling

To assess the functional interaction landscape of MYO1E in epithelial cells, we performed in situ proximity labelling using BirA*-fused MYO1E tail constructs in retinal pigment epithelial (RPE) cells. The MYO1E tail includes the PH domain in TH1, the proline-rich TH2 region, and the SH3 domain. RPE cell lines stably expressing BirA*-MYO1E tail (aa 692-1108), and mutant constructs lacking either the SH3 domain (ΔSH3) or with point mutations in the PH domain (ΔPH), were generated and validated for expression (Supplementary Figure 1B). Biotinylated proteins were enriched via streptavidin pulldown following overnight biotin labelling and analysed by mass spectrometry. Data from 3 independent experiments for each construct were compared against 6 BirA*-only control pulldowns.

Proteins with a 4.5-fold increase over control were considered significantly enriched. As shown in the heatmap in Figure 2A, a distinct set of MYO1E-associated proteins was identified, some of which overlapped with the MYO1F interactome. Volcano plot analysis (Figure 2B) revealed 45 high-confidence MYO1E-interacting proteins, including known actin-binding and membrane-associated proteins such as ITGA5 ^55^, FERMT2 (also known as kindlin-2)^56^, and CDC42EP1^57^. Disruption of the SH3 domain significantly altered the interactome, as illustrated in Figure 2C, indicating that this domain is critical for specific protein interactions. In contrast, mutation of the PH domain had very little effect on the interactome (Figure 2A).

**Figure 2:**
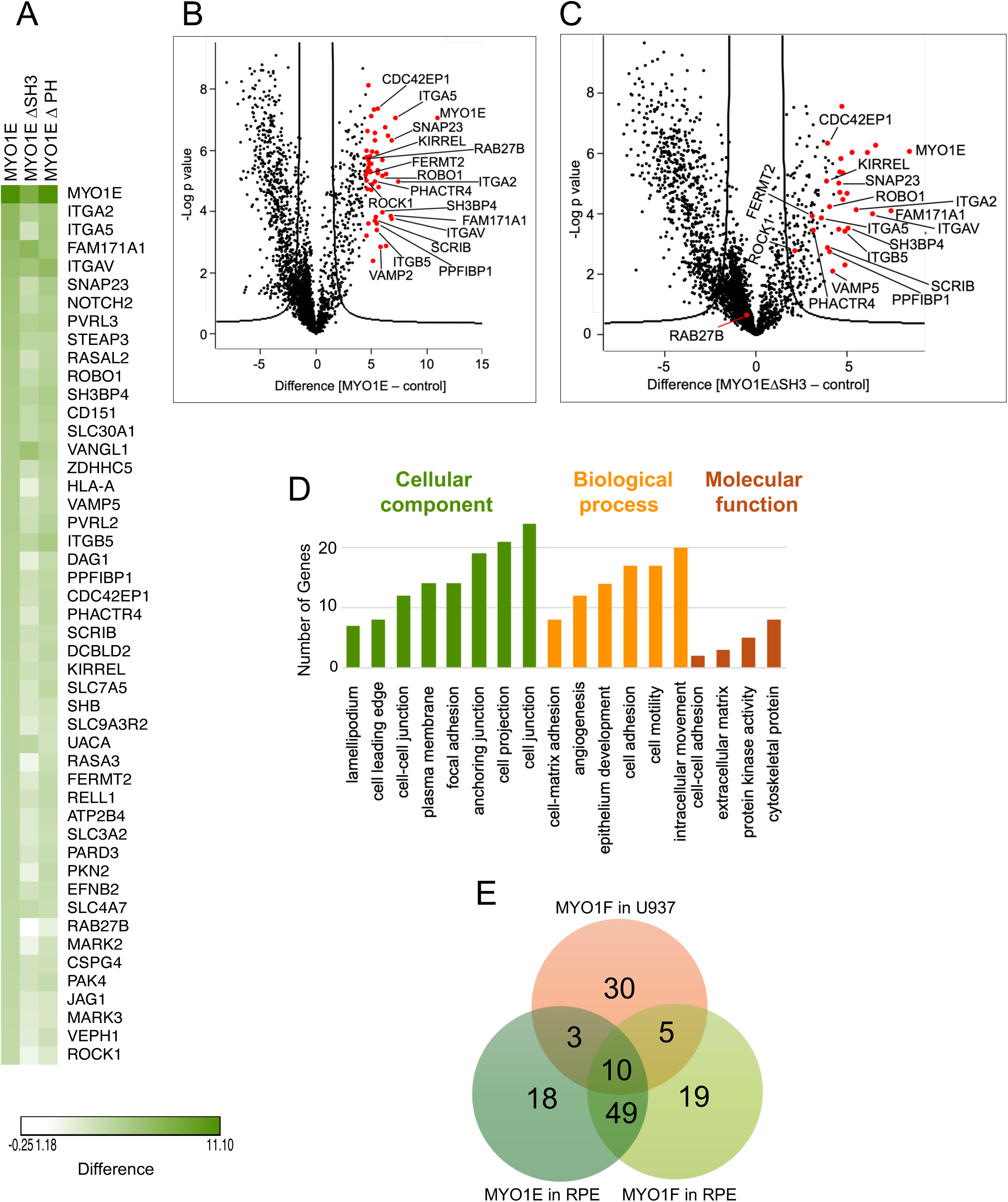
In situ proximity labelling reveals new binding partners for MYO1E. **A**. Heat map of abundance of protein interactions with MYO1E, MYO1EΔSH3 or MYO1EΔPH determined by BioID. Shown are protein interactions with a difference ≥ 4.5 over control. Data is shown from 3 independent experiments for MYO1E, and 3 independent experiments for the two mutants. **B**. Volcano plot of LFQ data from BirA*-MYO1E-tail over BirA* control from RPE-stable cell lines. The negative logarithmic *P* value was plotted against the t-test difference (n=3). The hyperbolic cutoff curve delimitates significantly enriched proteins from common hits (FDR of 0.01 and SO of 2). Significantly enriched (red dots) and other notable proteins are labelled by name. The complete list is given in supplementary data. **C**. Volcano plot of LFQ data from BirA*-MYO1E-tailΔSH3 over BirA* control. The negative logarithmic *P* value was plotted against the t-test difference (n=3). The hyperbolic cutoff curve delimitates significantly enriched proteins from common hits (FDR of 0.01 and SO of 2). **D**. Gene ontology (GO) classification of MYO1E interactome identified by in situ proximity labelling. GO cellular component, biological process and molecular function enrichment analyses were performed using ShinyGO software (Ge SX, Jung D and Yao R. Bioinformatics 36: 2628-2629, 2020). A term with P < 0.05 was considered to be significantly overrepresented. **E**. Venn diagram showing the relative numbers of significant interacting proteins detected for MYO1F in U937 cells, MYO1F in RPE cells and MYO1E in RPE cells in three independent BioID experiments. The overlap displays those proteins which were found in multiple constructs. The Venn diagram was made by using the web page https://bioinformatics.psb.urgent.be/webtools/venn/.

To further explore the potential functional implications of the MYO1E interactome, we performed Gene Ontology (GO) enrichment analysis. Proteins significantly enriched in the MYO1E interactome were associated with actin cytoskeleton organisation, membrane ruffling, and cell-substrate junction assembly (Figure 2D). This supports the notion that MYO1E plays a role in cell adhesion and cytoskeletal remodelling in epithelial cells, consistent with its known role in podosome function^28,21^.

Comparison of the interactomes of MYO1E in RPE cells with MYO1F in both U937 and RPE cells revealed overlapping and distinct protein networks (Figure 2E). A total of 10 proteins were shared among all three datasets, whereas 30 proteins were unique to MYO1F in U937 cells and 18 were specific to MYO1E in RPE cells and 19 for MYO1F in RPE cells. Interestingly, whereas only 3 proteins were shared between MYO1F in U937 and MYO1E in RPE cells this number was 49 when the proximity ligation experiments were performed for both myosins in RPE cells suggesting cell type -specific binding partners for MYO1F. These findings highlight both the conserved and divergent roles of long-tailed class I myosins across different cell types and suggest that MYO1E and MYO1F may perform overlapping yet distinct functions through differential binding partner selection.

### Validation of the CASS complex as a MYO1F-associated adaptor protein complex

To validate MYO1F-binding partners identified by proximity labelling, we performed co-immunoprecipitation (co-IP) assays with GFP-tagged MYO1F and candidate adaptor proteins in RPE cells. Among the most enriched hits in the MYO1F interactome were components of the CASS complex: CD2AP, ASAP1, SH3BP2 and SH3KBP1 (Figure 3A). These proteins are characterised by multiple SH3 domains, proline-rich regions, or PH domains (Figure 3B), suggesting the potential for multivalent interactions with MYO1F.

**Figure 3.**
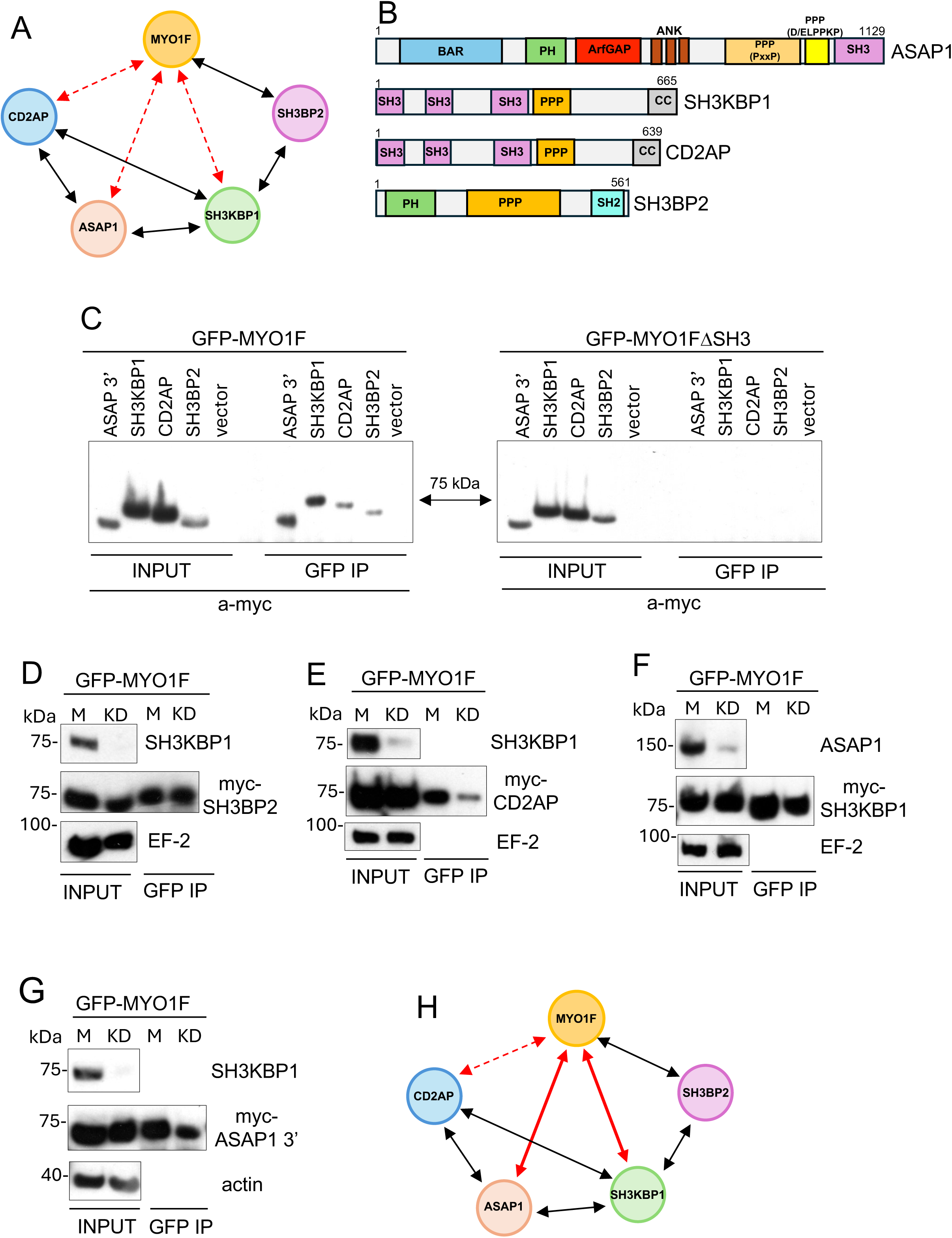
Verifying the CASS protein complex as part of the MYO1F interactome by immunoprecipitation. **A**. Network diagram of the CASS complex comprising <u>C</u>D2AP, <u>A</u>SAP1, <u>S</u>H3KBP1 and <u>S</u>H3BP2. Black arrows highlight known interactions and red dotted arrows suggested interactions from the BioID experiments. **B**. Schematic cartoon of ASAP1, SH3KBP1, CD2AP and SH3BP2 domain organisation. **C**. Immunoprecipitations were performed using GFP-antibodies from RPE cells stably expressing GFP-MYO1F (left panel) or GFP-MYO1FΔSH3 (right panel) and transiently transfected with myc-ASAP-3’ (aa 589 - aa1129), myc-SH3KBP1, myc-CD2AP, myc-SH3BP2 or empty myc-control vector. Input and immunoprecipitates were analysed by Western blotting with anti-myc antibodies. **D**. Immunoprecipitations were performed with GFP-antibodies from GFP-MYO1F expressing RPE cells either mock transfected (M) or transfected with siRNA oligonucleotides to SH3KBP1 (KD) before transfection with myc-SH3BP2. Immunoprecipitates were analysed by Western blotting with antibodies to endogenous SH3KBP1 and myc. EF2 was used as a loading control. **E**. GFP-MYO1F was immunoprecipitated from RPE cells either mock transfected (M) or transfected with siRNA oligonucleotides to SH3KBP1 (KD) before transfection with myc-CD2AP. Immunoprecipitates were analysed by Western blotting with antibodies to endogenous SH3KBP1 and myc. EF2 was used as a loading control. **F**. GFP-MYO1F was immunoprecipitated from RPE cells either mock transfected or transfected with siRNA oligonucleotides to ASAP1 before transfection with myc-SH3KBP1. Immunoprecipitates were analysed by Western blotting with antibodies to endogenous ASAP1 and myc. EF2 was used as a loading control. **G**. GFP-MYO1F was immunoprecipitated from RPE cells either mock transfected or transfected with siRNA oligonucleotides to SH3KBP1 before transfection with myc-ASAP3’. Immunoprecipitates were analysed by Western blotting with antibodies to endogenous SH3KBP1 or myc. Actin was used as a loading control. **H**. Interaction network diagram of the CASS-complex. Black arrows highlight known interactions and red solid lines confirmed direct interaction with MYO1F.

In GFP-pull down experiments from RPE cells expressing both GFP-MYO1F and Myc-tagged versions of each candidate, we observed co-precipitation of all four proteins with full-length MYO1F (Figure 3C, left panels). In contrast, deletion of the SH3 domain (MYO1FΔSH3) abolished binding of CD2AP, SH3KBP1, SH3BP2 as well as ASAP1 indicating that these interactions are dependent on the SH3 domain (Figure 3C, right panels).

To further dissect the architecture of the CASS complex, we performed siRNA knockdowns of SH3KBP1 and ASAP1 before pulling down GFP-MYO1F. Knockdown of SH3KBP1 had no effect on the binding of SH3BP2 to MYO1F indicating direct interaction (Figure 3D). However, knockdown of SH3KBP1 markedly reduced interaction of CD2AP with MYO1F (Figure 3E), indicating mostly indirect binding of CD2AP via SH3KBP1 to MYO1F. Conversely, knockdown of ASAP1 did not reduce SH3KBP1 co-precipitation with MYO1F (Figure 3F) and vice versa (Figure 3G), suggesting that both SH3KBP1 and ASAP1 can bind directly to MYO1F.

These results confirm the physical association of MYO1F with a multi-protein complex comprising CD2AP, ASAP1, SH3BP2 and SH3KBP1. The interaction network (Figure 3H) reflects both previously known interactions (shown by black arrows) and newly established interactions between MYO1F and ASAP1 as well as between MYO1F and SH3KBP1 (highlighted by red arrows). The SH3 domain of MYO1F enables multivalent binding to proline-rich regions found in all members of the CASS complex. Except for SH3BP2, each of these members also contains at least one SH3 domain, facilitating further network formation within the complex.

### In vivo experiments confirm the AlphaFold3-predicted binding interface between the proline-rich domain of SH3BP2 and the MYO1F-SH3 domain

To further analyse the molecular basis of the interaction between MYO1F and SH3KBP1 or ASAP1, we first generated GFP-tagged MYO1F tail constructs with deletions of 20, 45, and 57 amino acids at the C-terminus of the SH3 domain (Figure 4A). The SH3 domain of MYO1F comprises 58 amino acids and adopts a canonical fold characterized by five antiparallel β-strands forming a compact β-barrel structure and a short 3₁₀ helix. SH3 domains typically recognize proline-rich motifs with the consensus XPxXP, where X is a hydrophobic residue and x any amino acid. These motifs engage hydrophobic grooves within the SH3 domain via conserved proline residues, and binding specificity is often enhanced by a third, negatively charged pocket that interacts with flanking positively charged residues. Co-immunoprecipitation from transiently transfected RPE cells revealed that deletion of the terminal 20 amino acids markedly reduced MYO1F binding to SH3KBP1 and ASAP1 (Figure 4B). This region includes N1093 and Y1094, which are predicted by AlphaFold3 to contribute to the interaction with the proline-rich region of SH3BP2 (Figure 4D and F).

**Figure 4.**
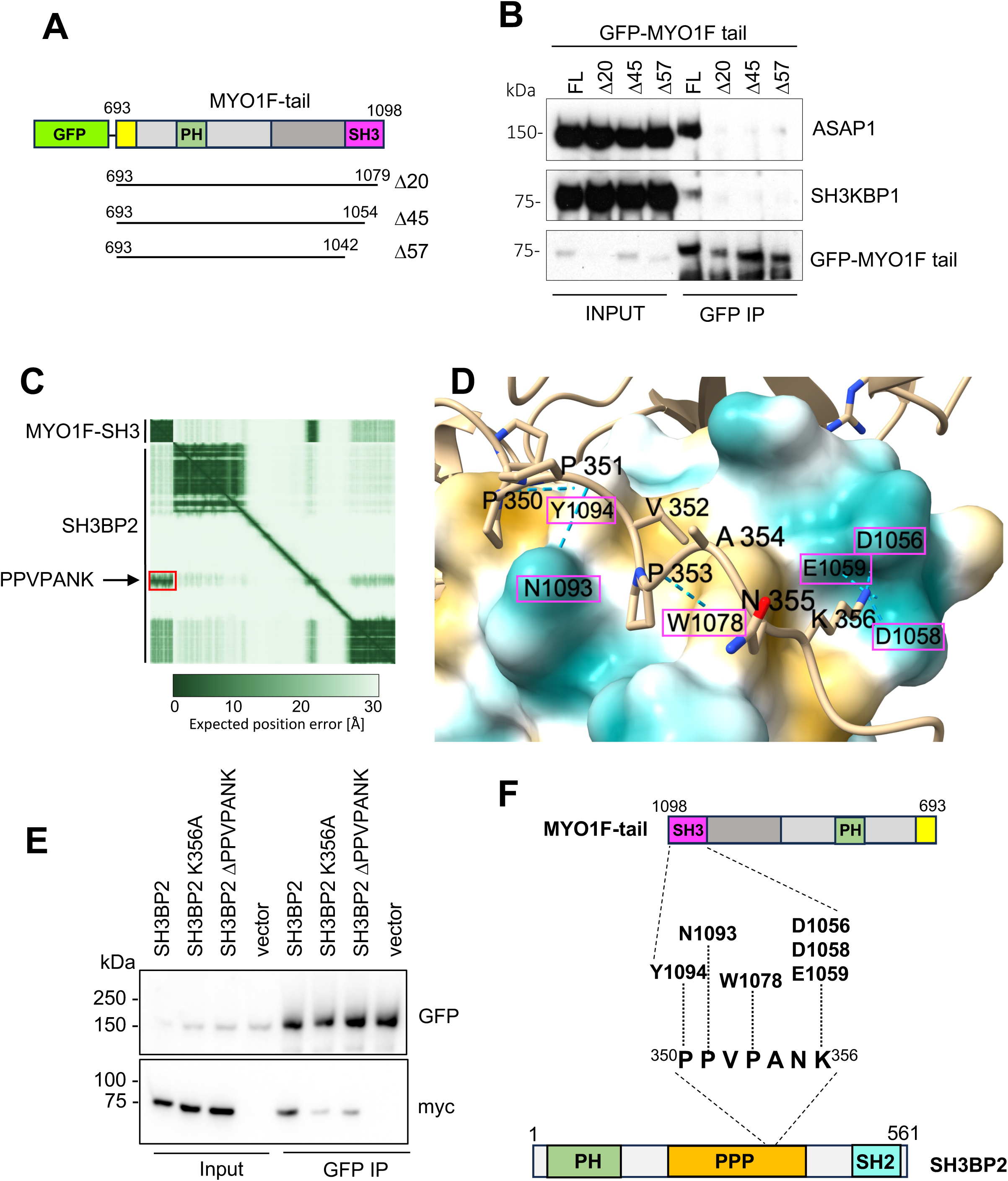
Characterisation of the molecular interaction between MYO1F SH3-domain and the SH3BP2 proline-rich domain. **A**. Schematic representation of the MYO1F tail domain highlighting the position of 20, 45 and 57 amino acids deletions at the C-terminus of the SH3-domain. **B**. Immunoprecipitations were performed from RPE cells transiently expressing either full-length GFP-MYO1F tail or C-terminal deletions of 20, 45 and 57 amino acids using antibodies to GFP. Immunoprecipitates were analysed by Western blotting with antibodies to endogenous ASAP1 or SH3KBP1. **C**. Alphafold3-predicted interaction between MYO1F-SH3 domain (residues 1042-1098) and full-length SH3BP2 (residues 1-651). The predicted aligned error (PAE) plot reveals a high confidence interaction between the proline-rich region of SH3BP2 (PPVPANK boxed in red) and the MYO1F-SH3 domain. The green gradient in the PAE plot indicates confidence levels, with darker regions representing higher prediction confidence. **D**. AlphaFold3 predicted structural model showing the interaction interface between the SH3BP2 PPVPANK motif (aa 350-356, shown as beige stick structure) and part of the MYO1F-SH3 domain (interacting amino acids highlighted by pink boxes). The PPVPANK motif is positioned within a hydrophobic region of the MYO1F-SH3 domain (highlighted in yellow). Predicted hydrogen bonding is shown by dashed blue lines. **E**. Immunoprecipitation from RPE cells stably expressing GFP-MYO1F and transiently expressing myc-tagged SH3BP2 constructs (wild-type, K356A point mutant, or a deletion mutant lacking the PPVPANK motif) using GFP-specific nanobodies. Immunoprecipitations were analysed by immunoblotting with antibodies against GFP and myc. **F.** Schematic summary of the predicted molecular interaction between SH3BP2 and the MYO1F SH3 domain, emphasizing the role of the PPVPANK motif in binding.

To further explore the SH3-domain–ligand interface, we used AlphaFold3 to model the interaction between the MYO1F SH3 domain (aa 1042–1098) and full-length SH3BP2 (aa 1–651). Alphafold3 modelling predicts a high-confidence interaction (iPTM=0.87), shown as a red boxed area on the predicted aligned error (PAE) plot, between the MYO1F SH3 domain and a short proline-rich motif in SH3BP2 (PPVPANK; aa 350– 356) (Figure 4C). The ipTM score provides an estimate of the predicted accuracy of the relative positioning between interacting protein domains, with values above 0.8 indicating high-confidence, high-accuracy predictions, whereas scores between 0.6 and 0.8 represent lower-confidence models that may be either correct or incorrect. The SH3BP2 proline-rich motif fits into a hydrophobic groove on the MYO1F SH3 domain, with multiple predicted hydrogen bonds stabilizing the interaction (Figure 4D and F). Structural modelling revealed that prolines P350 and P351 in SH3BP2 bind to residues Y1094 and N1093 (not in the hydrophobic groove) in the MYO1F SH3 domain, forming one hydrophobic groove, while P353 interacts with W1078 in a second hydrophobic pocket. In addition, the terminal lysine (K356) of the motif docks into a negatively charged specificity pocket formed by D1056, D1058, and E1059 (Figure 4D and F).

To validate this predicted binding interface in cells, we generated myc-tagged SH3BP2 constructs with either a point mutation in the predicted core lysine (K356A) or a deletion of the entire PPVPANK motif (ΔPPVPANK). These constructs were co-expressed with GFP-MYO1F in RPE cells, and GFP-based pull-downs were performed. Western blotting revealed that both the K356A substitution and the PPVPANK deletion significantly reduced the interaction with MYO1F, confirming the functional importance of this motif (Figure 4E). Interestingly, deleting the PPVPANK motif does not fully abolish the interaction with MYO1F, suggesting that the additional proline-rich sequences present within SH3BP2 may allow interaction with the SH3 domain.

Taken together, these findings provide structural and biochemical evidence that the MYO1F SH3 domain recognizes the PPVPANK motif in SH3BP2 through canonical SH3 domain interactions, involving two hydrophobic grooves and a specificity pocket. The PPVPANK motif is thus the principal determinant for high-affinity SH3-domain-mediated binding (Figure 4F).

### Identification of multiple proline-rich motifs in ASAP1 that mediate binding to the MYO1F SH3 domain

To further dissect the molecular interface between MYO1F and ASAP1, we used AlphaFold3 to model the interaction between the MYO1F SH3 domain and the C-terminal half of ASAP1 (residues 589–1129). Structural predictions revealed several proline-rich motifs (PRMs) in ASAP1 that could potentially interact with the MYO1F SH3 domain, with distinct confidence scores indicated by ipTM values (Figure 5A–B). In the D/ELPPKP domain several PPKP motifs between residues 937-993 (region A in constructs 1 and 2) have been identified, which give a lower-confidence ipTM score of 0.66 and 0.69 (Figure 5B). Consistent with the lower ipTM score, co-immunoprecipitation assays using GFP-MYO1F and myc-tagged ASAP1 truncation constructs without these motifs showed only slight reduction in MYO1F binding (construct 3, residue 589-937), suggesting that these predicted motifs are not essential for the observed interaction (Figure 5C).

**Figure 5.**
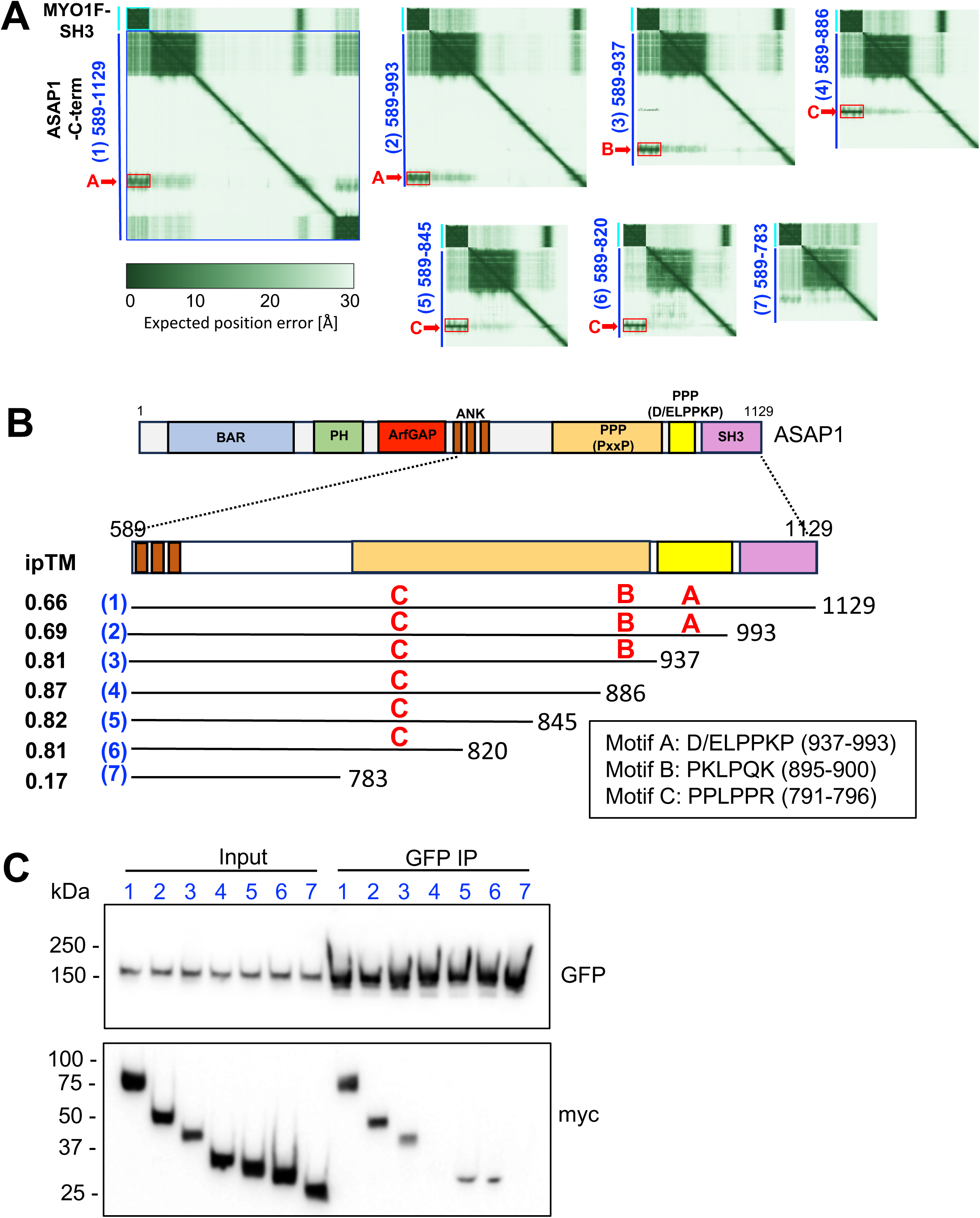
Experimental verification of predicted MYO1F-ASAP1 interaction. **A**. AlphaFold3-predicted interaction of MYO1F with the C-terminus of ASAP1 (residues 589–1129) and six truncation constructs. Each PAE plot highlights intramolecular confidence of interaction, where darker green represents higher confidence. Red boxes denote predicted binding regions aligning with conserved proline-rich motifs (A– C). Constructs are labelled (1) through (7) with reducing lengths. **B**. Schematic illustration of full-length ASAP1, illustrating key domains: BAR (Bin-Amphiphysin-Rvs), PH (Pleckstrin Homology), ANK (ankyrin repeats), PPP (Proline-Rich) (PxxP) or (D/ELPPKP repeats), and SH3. Enlarged C-terminus of ASAP1 (residue 589-1129) with overview of ASAP1 truncation constructs (1-7 in blue) tested for MYO1F binding underneath. The proline-rich domain of ASAP1 contains several proline-rich motifs (PRM) (A-C) that are predicted by AlphaFold3 to bind the MYO1F SH3-domain. RPMs are labelled A–C in red with their predicted ipTM score on the left-hand side. **C**. Immunoprecipitation from RPE cells stably expressing GFP-MYO1F and transiently expressing myc-tagged ASAP1 truncation constructs (1–7). GFP pull-downs were performed using GFP-specific nanobodies. Input and immunoprecipitated fractions were analysed by Western blotting with anti-myc and anti-GFP antibodies.

To map the minimal region of ASAP1 required for MYO1F interaction, further C-terminal truncations in combination with AlphaFold-prediction were generated between residues 589–937 and assessed for MYO1F binding (Figure 5C). While deletion of the class II motif (PKLPQK) at position (895-900), does not completely abolish ASAP1-MYO1F interaction (see construct 5 and 6), deletion of residues 784– 820 (construct 7) abrogated MYO1F binding, suggesting that a PRM within this region mediates binding. AlphaFold3 predicted a class II PPLPPR motif (residues 791-796, motif C) in this segment with an ipTM of 0.87. Supporting this prediction, deletion of this region abolished binding, while its inclusion in construct 5 and 6 retained MYO1F interaction. Construct 4 appears to be an anomaly, as binding to MYO1F is abolished despite retaining the putative binding site in motif C. This observation may suggest that intramolecular folding or structural context also influence motif accessibility or binding affinity.

In summary, AlphaFold3 predictions together with experimental truncation mapping highlight that the MYO1F SH3 domain has the potential to interact with multiple PRMs in the C-terminal region of ASAP1. These results suggest that MYO1F exhibits binding plasticity across several partially redundant PRMs potentially allowing motor multimerization.

### Subcellular localisation of MYO1F and its adaptor proteins in THP-1 macrophages

We next examined the subcellular localisation of endogenous MYO1F and its validated direct binding partners, ASAP1, SH3KBP1, and SH3BP2, in myeloid cells using high-resolution immunofluorescence microscopy in phorbol 12-myristate 13-acetate (PMA)-differentiated THP-1 macrophages. Confocal microscopy of cells stained with anti-MYO1F antibodies and phalloidin revealed that MYO1F localises to discrete actin-rich puncta at the ventral cell surface, consistent with the organisation of podosomes (Figure 6A)^22^. These MYO1F-positive puncta were spatially restricted to the basal cortex, as shown in the z-plane proximal to the coverslip. Our observations confirm the previously reported identification of MYO1F as a podosome component by mass spectroscopy^20^. Furthermore, both ASAP1 and MYO1E, the closest mammalian homologue of MYO1F, have been localised at podosomes^21,58^. While ASAP1 has been implicated in podosome formation and turnover through interactions with proteins such as GEFH1^59^ and FAK^58^, MYO1E has been suggested to regulate actin polymerisation and membrane-cytoskeleton coupling in a phosphatidylinositol (3,4,5)-trisphosphate (PI (3,4,5) P3)-dependent manner ^28^.

**Figure 6.**
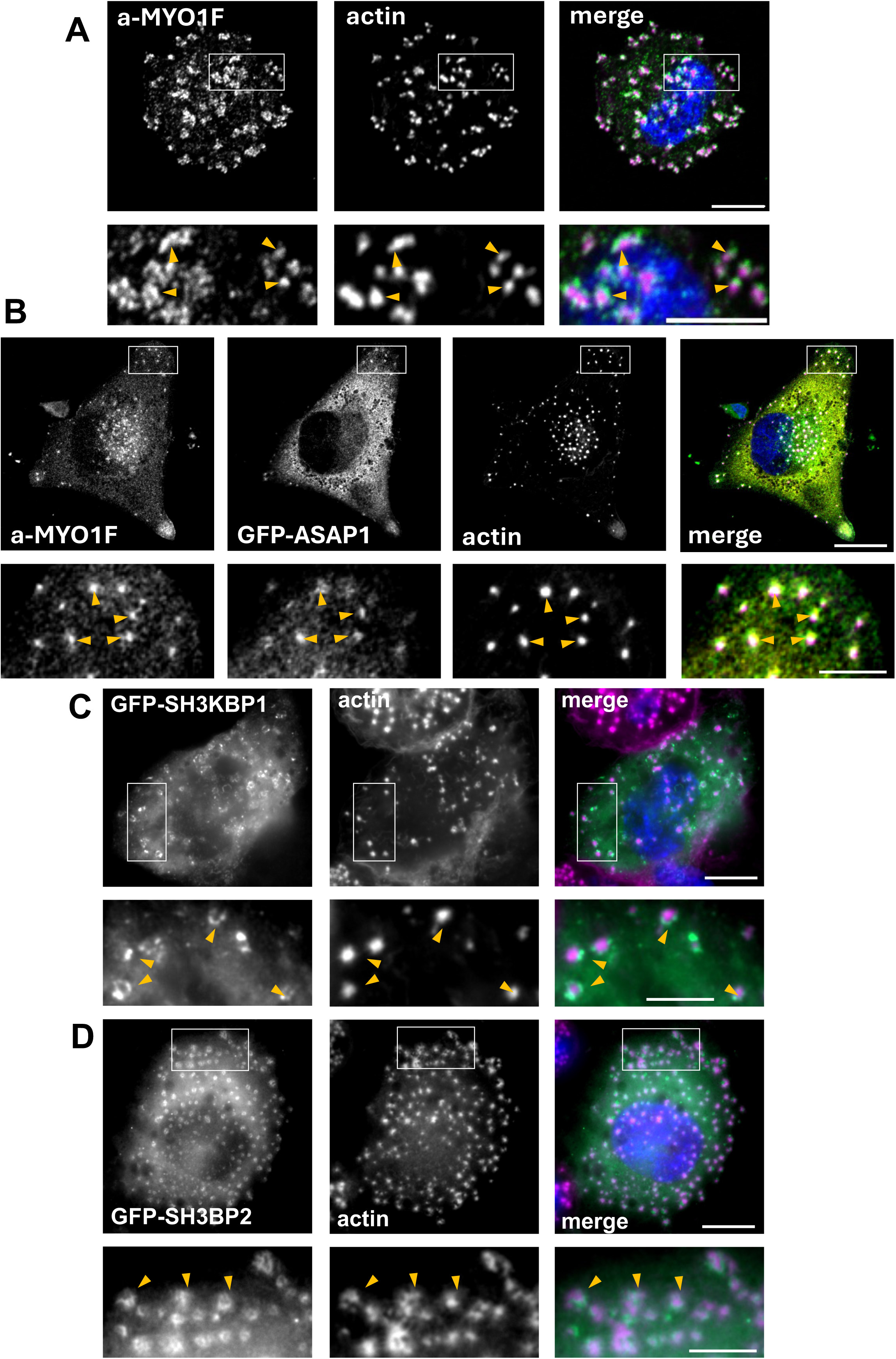
MYO1F and ASAP1 colocalise in ventral podosomes in macrophages. **A.** THP-1 cells treated with PMA to induce differentiation into adherent macrophage-like cells were stained in immunofluorescence with antibodies to MYO1F and double labelled with fluorescently labelled phalloidin to visualise actin filaments. In the merge image on the righthand panel actin is labelled in magenta and MYO1F in green. Confocal images were taken at the dorsal layer of the cell close to the coverslip. **B.** THP-1 cells were transiently transfected with GFP-ASAP1and labelled with antibodies to GFP and MYO1F and stained with fluorescently labelled phalloidin to visualise actin filaments. The merge image on the righthand panel shows actin in magenta, GFP-ASAP1 in green and a-MYO1F in cyan. Confocal images were taken at the dorsal layer of the cell close to the coverslip in **A** and **B**. **C** and **D.** THP-1 cells transiently expressing GFP-SH3KBP1 (**C)** or GFP-SH3BP2 (**D)** and labelled with antibodies to GFP and fluorescently labelled phalloidin to visualise actin filaments. The merge image on the righthand panel shows actin in magenta and GFP-SH3KBP1 or GFP-SH3BP2 in green. Widefield images focused on the dorsal part of the cell are shown. Confocal images were taken at the dorsal layer of the cell close to the coverslip. White boxes in **A, B**, **C and D** indicate areas enlarged in the picture below. Orange arrowheads indicate colocalization. Scale bar 10 µm.

To further compare the spatial distribution of MYO1F and its binding partner ASAP1, we co-stained THP-1 macrophages transiently expressing GFP-ASAP1 with anti-MYO1F antibodies and phalloidin. Confocal imaging revealed colocalisation of GFP-ASAP1 and endogenous MYO1F at ventral actin puncta (Figure 6B), indicating that both MYO1F and ASAP1 are recruited to podosomes.

To assess whether the adaptor proteins SH3KBP1 and SH3BP2 colocalise with MYO1F at podosomal structures, we transiently expressed GFP-tagged constructs in THP-1 macrophages and performed widefield imaging focused on the ventral plane. Both GFP-SH3KBP1 and GFP-SH3BP2 exhibited punctate localisation patterns similar to MYO1F that overlapped with phalloidin-labelled F-actin (Figure 6C–D), indicating recruitment to podosome-like structures. Enlarged views highlighted the close spatial alignment of these adaptors with actin foci (arrowheads), supporting their proposed localisation at podosomes.

These observations confirm that MYO1F and members of the CASS complex (ASAP1, SH3KBP1, SH3BP2) are recruited to actin-rich podosomes in macrophages, supporting a functional role for this protein complex in cytoskeletal organisation and membrane-cytoskeleton coupling.

### MYO1F and ASAP1 colocalise in podosomes in primary mouse microglia and iPSC-derived human microglia

Given the upregulation of MYO1F expression in microglia within both murine models of neurodegeneration and AD patient tissue^15,14,18,16^, we examined the subcellular localisation of endogenous MYO1F and its adaptor protein ASAP1 in both primary mouse microglia and human induced pluripotent stem cell (hiPSC)-derived microglia to assess whether recruitment to podosomes is conserved across different myeloid cell types.

Immunofluorescence microscopy of primary mouse microglia revealed that MYO1F localises to discrete phalloidin-labelled F-actin structures at the ventral surface of the cell suggesting the recruitment of MYO1F to podosomes in brain-derived microglia (Figure 7A).

**Figure 7:**
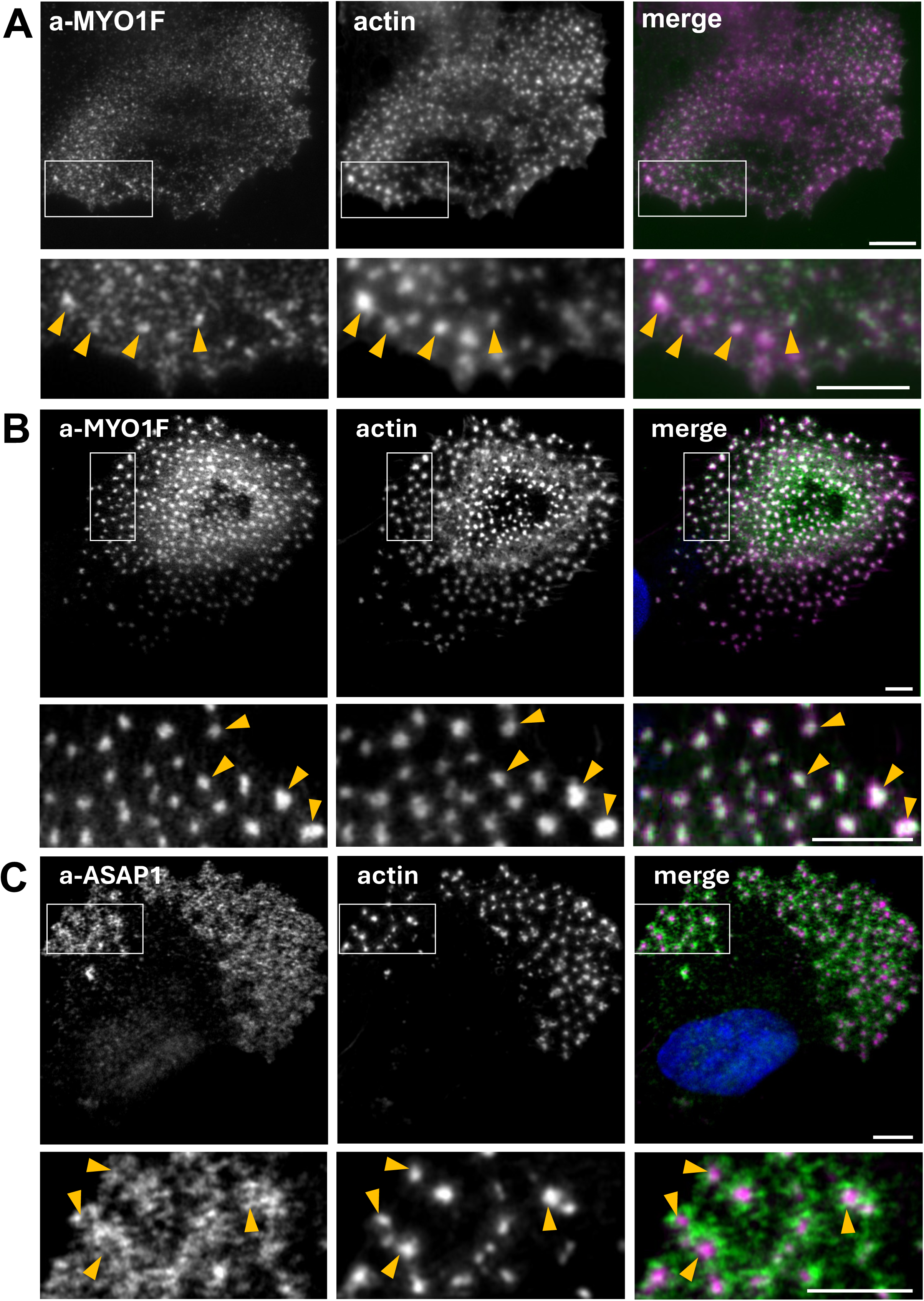
MYO1F and ASAP1 colocalise in podosomes in primary mouse microglia and iPSC-derived human microglia. **A**. Primary mouse microglia on coverslips were stained with antibodies to MYO1F and labelled with phalloidin to visualise actin filaments. Orange arrowheads indicate colocalization. Widefield images focused on the dorsal part of the cell are shown. White boxes indicate areas enlarged in the picture below. Scale bar 5 μm. **B**. and **C**. Human iPSC-derived microglia were stained with antibodies to MYO1F (**B**) or ASAP1 (**C**) and labelled with phalloidin to label actin filaments. The merge image on the righthand panel shows actin in magenta, MYO1F or ASAP1 in green and nucleus in blue. Confocal images were taken at the dorsal layer of the cell close to the coverslip. White boxes in **A, B** and **C** indicate areas enlarged in the picture below. Orange arrowheads indicate colocalization. Scale bar 5 μm.

To determine whether these observations extend to human microglia, we assessed the localisation of MYO1F and ASAP1 in human iPSC-derived microglia. Confocal imaging revealed that both proteins displayed punctate distributions at the basal cortex, colocalising with F-actin-rich domains (Figure 7B–C). These structures were morphologically consistent with podosomes observed in peripheral myeloid cells and primary mouse microglia. Merged images confirmed substantial colocalisation of MYO1F and ASAP1 with the actin cytoskeleton in ventral domains, supporting their recruitment to podosomes.

These findings confirm that MYO1F and ASAP1 localise to podosomes not only in macrophages but also in both mouse and human microglia. Given the known involvement of podosomes in adhesion and matrix remodeling^23^, and the reported transcriptional upregulation of MYO1F in neuroinflammatory states associated with neurodegenerative disease^14,15^, the conserved podosomal recruitment of MYO1F and its adaptor protein ASAP1 supports a role for this complex in cytoskeletal remodeling and immune function within the central nervous system.

### MYO1F recruitment to the phagocytic cup requires motor activity and adaptor proteins

To investigate the mechanisms underlying MYO1F localization during phagocytosis, we examined the recruitment of MYO1F and its known adaptor proteins to phagocytic cups. Both MYO1F and its paralogue MYO1E are enriched at the phagocytic cup, where they regulate cortical tension and promote phagosome closure^31,33,60,32^. Interestingly, podosome-like structures, called phagocytic podosomes, can also be observed at the ventral edge of the phagocytic cup^29^. These structures may function as adhesion sites that generate traction for particle internalization.

To determine which MYO1F-associated adaptor proteins are recruited to the phagocytic cup, we performed three-color confocal imaging of RAW264.7 macrophages transfected with Lifeact-mApple and GFP-tagged constructs of MYO1F binding partners (Figure 8A). Following the addition of Alexa647-labelled BSA-coated beads to stimulate phagocytosis, we assessed the accumulation of each protein at the actin-rich phagocytic cup (Figure 8B). In this assay we included the analysis of utrophin (UTRN), another putative MYO1F adaptor protein (Figure 1). UTRN is a very large protein, and so we used a shorter form, only containing the actin binding calponin homology domains (CH1-CH2). Quantifying the recruitment of the putative binding partners we observed that the UTRN CH1-CH2 domain (0.4±0.44), CD2AP (0.32±0.41) and SH3BP2 (0.49±0.45) accumulated almost equally at the phagocytic cup, while SH3KBP1 showed lower enrichment (0.14 ± 0.34) (Figure 8B). ASAP1 was not included in this assay as it is not expressed in RAW macrophages.

**Figure 8:**
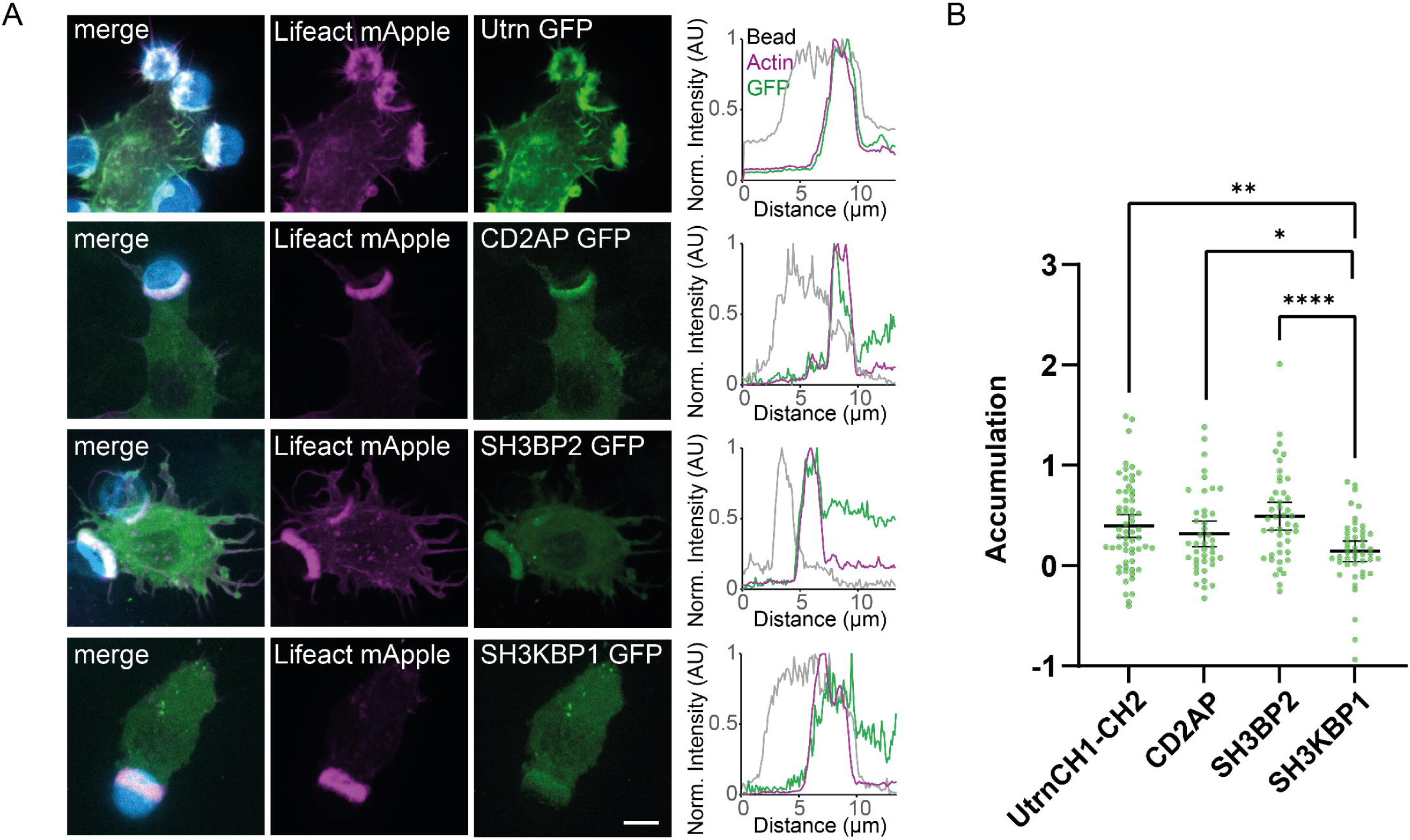
Recruitment of MYO1F binding partners at the phagocytic cup. **A**. Fluorescence images of RAW 246.7 cells, expressing Lifeact (magenta) and the binding partner of interest (green), engulfing Alexa647 labelled beads (blue). Scale bar 5µm. Line profiles across the dashed line (white, merge) show the normalised fluorescence intensity of the bead, actin and the protein of interest (right). **B**. Dot plot of accumulation values comparing the proteins UTRN CH1-CH2 (n=60), CD2AP (n=42), SH3BP2 (n=44) and SH3KBP1 (n=46). Data shown was collected from 3 independent experiments. Bars represent the mean ± 95% confidence interval. P values are represented as p<0.0001:****, p<:0.01:**, and p<0.05: *.

To define the structural features required for MYO1F localization during phagocytosis, we next compared the recruitment of full-length MYO1F with mutant constructs lacking specific domains using the same assay (Figure 9). Full-length MYO1F-GFP displayed robust enrichment at the phagocytic cup (0.89 ± 0.79) (Figure 9A and 9B) consistent with a role in linking membrane dynamics to cortical actin during particle engulfment ^32,31^. In contrast, mutation of the pleckstrin homology domain (ΔPH) or deletion of the SH3 domain (ΔSH3) led to significantly reduced accumulation (0.02 ± 0.20 and 0.16 ± 0.29, respectively). Expression of the tail-only construct, MYO1F-tail, which lacks the motor domain, showed no enrichment at the phagocytic cup (−0.16 ± 0.31) (Figure 9A and B).

**Figure 9:**
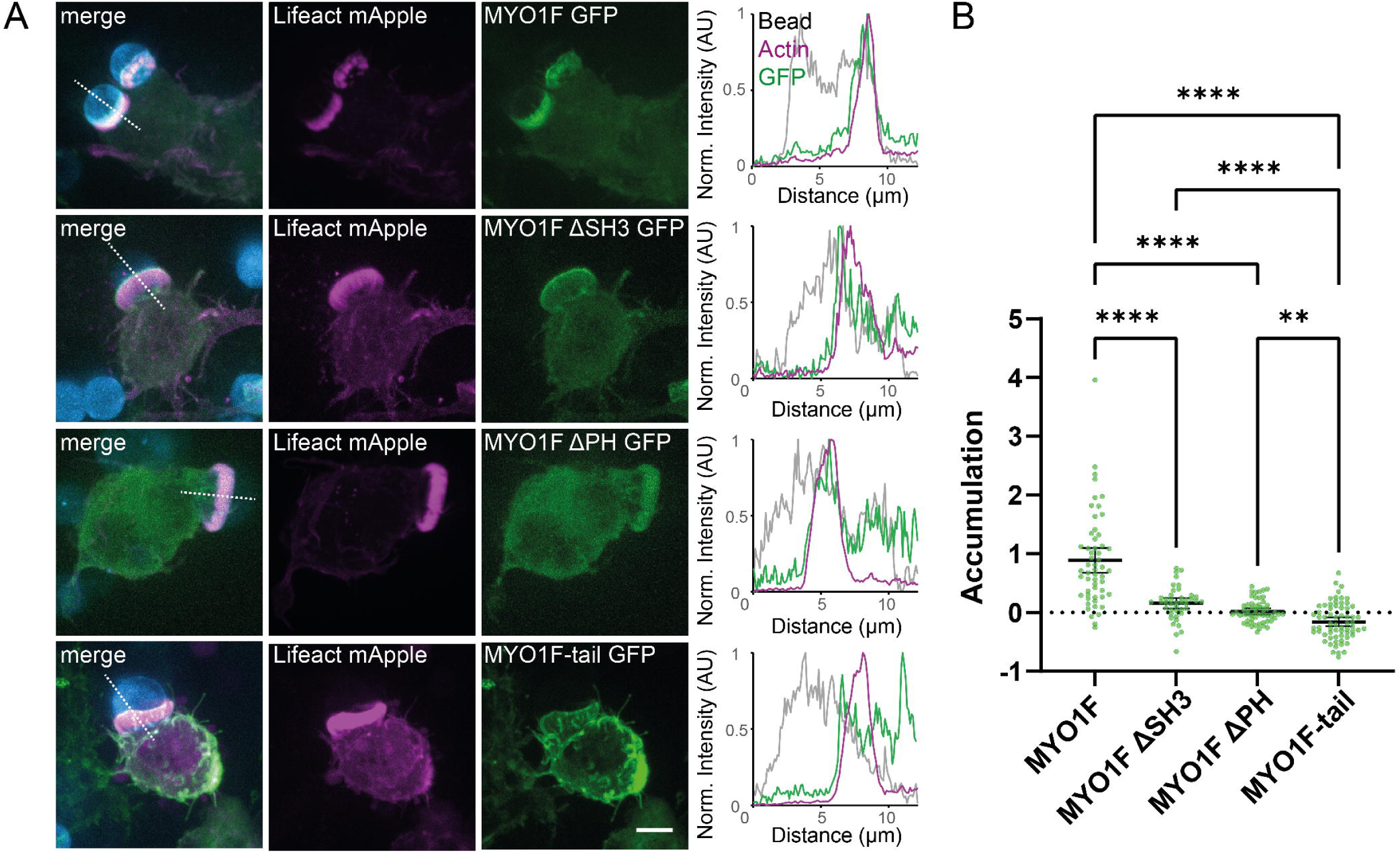
MYO1F recruitment to phagocytic cups requires the SH3 and PH domain. **A**. Fluorescence images of RAW 246.7 cells expressing Lifeact mApple (magenta) and either MYO1F, MYO1F ΔSH3, MYO1FΔPH or MYO1F-tail GFP (green). The cells are shown engulfing Alexa647 labelled beads (blue). Scale bar 5µm. Line profiles across the dashed line (white, merge) on the right show the normalised fluorescence intensity of the bead, actin and the protein of interest (right). **B**. Dot plot of accumulation values comparing cells expressing MYO1F (n=55) with MYO1F ΔSH3 (n=42), MYO1F ΔPH (n=56) and the tail of MYO1F (n=65). Data shown was collected from 3 independent experiments. Bars represent the mean ± 95% confidence interval. P values are represented as p<0.0001: ****, p<0.001:***, p<0.01:**.

In summary, these findings demonstrate that MYO1F recruitment to the phagocytic cup requires intact motor activity, membrane association via the PH domain, and adaptor protein binding via the SH3 domain. Together, these findings provide mechanistic support for the role of MYO1F in bridging membrane-cytoskeleton interactions during phagocytosis and underscore the functional convergence of podosome and phagosome biology.

## DISCUSSION

In this study we have used in situ proximity labelling-based proteomics to identify the interaction network of the two human long-tailed myosins of class I, MYO1E and MYO1F. We determined the interactome for both myosins using cell types in which they are endogenously expressed at high levels; RPE cells for MYO1E and both RPE and U937 cells for MYO1F. Using a combination of co-immunoprecipitation, mutagenesis, structural modelling and cell biological assays we characterised the functional CASS complex of CD2AP, ASAP1, SH3KBP1 and SH3BP2 that interacts with the C-terminal SH3-domain of MYO1F. In addition, we identified a number of membrane-localized proteins that depend on the intact MYO1F PH domain for binding, including NUMB, ADD3, PECAM1, GOLGA8R and GPR124.

Together, our proximity labelling and domain deletion strategies reveal two distinct functional modules within the MYO1F interactome: (1) a SH3-domain-dependent adaptor protein complex (CASS), involved in cytoskeletal organization and adaptor scaffolding, and (2) a PH-domain-associated set of membrane-associated proteins. These modules likely enable MYO1F to form a dynamic link between the plasma membrane and the actin cytoskeleton during processes such as podosome formation and phagocytosis.

Comparison of the interactomes revealed limited overlap between MYO1F in U937 and MYO1E in RPE cells reflecting their divergent tissue distributions and distinct functions. MYO1F expression is largely restricted to immune cells^12,13^, as confirmed by immunoblotting (Figure 1A), while MYO1E is broadly expressed across a variety of tissues including kidney, lung, intestine and immune cells^9,11^. In neutrophils, where both proteins are present at differing levels, MYO1E and MYO1F exhibit non-redundant roles. MYO1E deficiency caused increased rolling velocity, decreased adhesion, aberrant crawling, and strongly reduced transmigration through the capillary wall^61^. The highly expressed MYO1F is important for neutrophil migration as cells from the MYO1F KO mouse show stronger adhesion to integrin ligands consequently reducing cell motility under static conditions in vitro^12^. Finally, in mast cells MYO1F appears to regulate cortical actin filament dynamics affecting cell adhesion, migration and degranulation, a process that involves the adaptor protein SH3BP2, which we identified via in situ proximity labelling and verified using mutagenesis guided by AlphaFold-based structural predictions^42^. The display of these functional differences between MYO1E and F, when they are expressed in the same cell type, is supported by the only partially overlapping interactome. The limited sequence conservation in the tail region of only 55% identity does allow for distinct adaptor protein binding.

It has previously been demonstrated that the TH2 domain of MYO1E is essential for its podosomal localization, that the TH1 domain promotes its membrane enrichment and ventral localization, and that deletion of the SH3 domain does not impair its recruitment to podosomes^21,28^. While it was shown that the SH3 domain is dispensable for MYO1E, corresponding data for MYO1F were not presented. In contrast, our study reveals that the SH3 domain of MYO1F is essential for binding multiple podosome-resident adaptor proteins, including ASAP1, SH3KBP1, and SH3BP2. These interactions suggest that the SH3 domain of MYO1F is not crucial for recruitment but important for its functional engagement at podosomes and phagocytic structures. Notably, these binding partners are not shared with MYO1E, highlighting a mechanistic divergence from MYO1E and distinct recruitment strategies among long-tailed myosins of class I.

To date most functional studies on MYO1F have focused on peripheral immune cells, however emerging evidence indicates that MYO1F is also expressed in brain-resident microglia, the principal innate immune cells of the central nervous system. Although the precise molecular functions of MYO1F in microglia remain to be elucidated, our immunofluorescence studies in both primary mouse microglia and human iPSC-derived microglia reveal that MYO1F localises to ventral actin-rich structures resembling podosomes. These specialised adhesion sites are known to regulate cell-substrate interactions, matrix degradation, and mechanotransduction. The recruitment of MYO1F to these domains suggests that it may contribute to microglial adhesion, migration, and possibly cytokine secretion during neuroimmune responses.

There is gathering evidence for the dysregulation of peripheral and innate immune systems in neurodegenerative pathologies, including a skewed susceptibility of myeloid and microglial cells to inflammatory activation^62,63^. Given the selective upregulation of MYO1F in activated microglia in the context of neurodegenerative disorders, MYO1F may represent a novel therapeutic target, which warrants further investigations. Importantly, myosin motors are considered druggable proteins, exemplified by the development of small-molecule modulators for several myosin classes. Finally, the SH3KBP1-CD2AP complex binding to MYO1F has also been linked to neurodegeneration, particularly AD. CD2AP is an adaptor protein involved in actin, and membrane trafficking and genetic variants have been associated with increased risk for late-onset AD^64,65^. SH3KBP1 (also known as CIN85) is a scaffold protein also involved in endocytosis and receptor tyrosine kinase trafficking^66^. While less studied than CD2AP in neurodegeneration, emerging data implicate SH3KBP1 in neuroinflammatory pathways. The selective expression and disease-associated regulation of MYO1F thus highlights its potential as a pharmacological target for regulating microglial activation and attenuating neuroinflammation in conditions such as AD or FTD/ALS.

## ACKNOWLEDGEMENTS

We thank Dr. Robin Antrobus for help and advice with the proteomics and Wing-Hei Au for optimising the hiPSC-derived microglia differentiation protocol. This work was supported in the Buss laboratory by a program grant from the Medical Research Council (MR/S007776/1) and a studentship to EP co-funded by the Harding Distinguished Postgraduate programme and the School of Clinical Medicine, University of Cambridge doctoral training programme in Medical Research. CIMR microscopy is supported by equipment grants from the Wellcome Trust (108415) and the Medical Research Council (MR/Y002172/1). Work in the AL laboratory is supported by a UKRI Medical Research Council Senior Clinical Fellowship (MR/X006867/1) and funding in the GMG laboratory was provided by the Wellcome Trust (100140) and a Henry Wellcome Fellowship to AML (215899).

## SUPPLEMENTARY FIGURE LEGENDS

**Supplementary Figure 1: Verification of BirA-MYO1F or E expression by immunoblotting. A.** Analysis of U937 and RPE cells stably expressing myc-BirA*MYO1F tail, myc-BirA*MYO1F tailΔSH3 or myc-BirA*MYO1F tailΔPH or **B**. myc-BirA*MYO1E tail, myc-BirA*MYO1E tailΔSH3 or myc-BirA*MYO1E tailΔPH by immunoblotting using a myc antibody.

**Supplementary figure 2: Microglia marker proteins IBA1, P2Y12 and TMEM119 are expressed in iPSC-derived human microglia.** Human hiPSC-derived microglia were stained with antibodies to MYO1F and IBA1, P2Y12 or TMEM119 and labelled with phalloidin to label actin filaments. The merge image on the righthand panel shows actin in magenta, MYO1F in green, IBA1, P2Y12 or TMEM119 in yellow and nucleus in blue. Widefield images were taken at the dorsal layer of the cell close to the coverslip. Scale bar 5 μm.

**Supplementary Figure 3: Image analysis of phagocytosis assay.** Each fluorescence channel was first normalised to the maximum pixel intensity. Otsu thresholding was then calculated for each fluorescence channel to generate the protein of interest mask as well as the bead mask. To identify areas of high actin accumulation (the actin ring), the Otsu threshold for the actin channel was multiplied by two to generate a high actin mask. To compare the intensity of the protein of interest inside versus outside of the actin ring, two additional masks were generated, an inside actin mask and an outside actin mask. The inside actin mask was generated by multiplying the bead mask with the actin ring mask to ensure that only high actin structures engulfing a bead were included. The outside actin mask was created by subtracting the inside actin mask from the protein of interest mask. The average intensity of the protein of interest was then compared using an accumulation value which divided the average intensity of the protein of interest inside the actin ring by the average intensity outside the actin ring. The resulting value was then subtracted by one.

**Supplementary Table 1: MYO1F BioID Data.** Gene IDs, difference over control and p-value scores are shown.

**Supplementary Table 2: MYO1E BioID Data.** Gene IDs, difference over control and p-value scores are shown.

## MATERIAL AND METHODS

### Antibodies

The following antibodies were used: MYO1F (Santa Cruz Biotechnology, sc-376534), MYO1E (Atlas Antibodies, HPA023886), actin (Sigma, A2066), IBA1 (Wako, 019-19741), P2Y12 (Biolegend), GFP (Invitrogen, A11122), myc (Millipore, 05-724), SH3KBP1 (Santa Cruz Biotechnology, sc-166862), ASAP1 (Santa Cruz Biotechnology, sc-374410), EF2 (Santa Cruz Biotechnology, sc-166415), rabbit anti BSA (MPBiological 8651111). All secondary antibodies for immunofluorescence were purchased from Thermo Fisher Scientific.

### Plasmids

The MYO1F and CD2AP cDNAs were amplified by PCR from THP-1-derived cDNA, while the MYO1E tail domain and ASAP1 were amplified from RPE cDNA. SH3BP2 was amplified from HeLa cDNA, and SH3KBP1 (isoform 1) from U937 cDNA. Full-length or tail constructs were generated by PCR and subcloned into pEGFP-C, pCMV-myc, myc-BirA* pLXIN2, GFP-pLXIN2, and pHRSIN-GFP vectors. Site-directed mutagenesis was used to generate point mutations in the pleckstrin homology (PH) domains of MYO1F (K770A and R780A) and MYO1E (K722A and R782A). The pHRSIN-GFP vector was generated by replacing the mCherry sequence and multiple cloning site (MCS) from the PHRSIN-JJ vector (a kind gift from John James at Warwick University) with GFP and the MCS from pLXIN2, using XhoI and NotI restriction sites.

### Cell Culture

RPE (hTERT RPE-1; ATCC® CRL-4000™) cells were grown in 1:1 DMEM/F12-HAM (Sigma) supplemented with 10% fetal bovine serum (FBS) (Sigma), 2 mM L-glutamine (Sigma), 100 U/ml penicillin and 100 μg/ml streptomycin (Sigma). U937 cells were maintained in RPMI-1640 (Sigma) supplemented with 10% FBS, 2 mM L-glutamine, 100 U/ml penicillin and 100 μg/ml streptomycin. THP-1 cells were cultured in RPMI-1640 (Sigma) supplemented with 50µM beta mercaptoethanol, 10% FBS, 100U/ml penicillin and 100 μg/ml streptomycin. Differentiation of THP-1 cells into macrophage-like cells was induced by treatment with 50 ng/ml phorbol 12-myristate 13-acetate (PMA; Sigma P8139) for 24 hours, followed by a 48-hour recovery period in standard THP-1 medium. HEK293T and Phoenix cells (Phoenix-AMPHO; ATCC® CRL-3213™) were cultured in DMEM with GlutaMAX (Thermo Fisher Scientific) containing 10% FBS, 100 U/ml penicillin and 100 μg/ml streptomycin. Finally, RAW 246.7 macrophages were cultured in DMEM (Sigma, D5030) supplemented with 10% FBS and grown at 37°C, 10%CO_2_ and split every 2-3 days.

Stable RPE and U937 cell lines were generated using the Phoenix retroviral expression system with the pLXIN retroviral packaging vector. Virus was generated by transfection of Phoenix HEK293T cells with the pLXIN2 vector. The virus was harvested after 48h and used to infect target cells before selection with 0.5 mg/ml G418 (Gibco).

Transient transfections of RPE cells were performed using Fugene 6 (Promega) for immunofluorescence or PEI (Polysciences) for immunoprecipitations according to the manufacturer’s instructions.

Transient transfections of THP-1 cells were performed by lentiviral transduction using Lipofectamine 2000 (Thermo Fisher Scientific). Virus was generated by transfection of HEK293T cells with pHRSIN GFP and the packaging vectors gagpol and VSVG. After 48h virus was harvested and used to infect THP-1 cells.

Knockdowns were performed by transfection of ON-TARGETplus SMARTpool siRNA oligonucleotides into RPE cells using Oligofectamine (Thermo Fisher Scientific). To ensure efficient knockdown transfections were performed on days 1 and 3.

### Mouse microglia isolation

Primary microglia were isolated from mouse pups P1 to P3. Brains were extracted into Hanks’ Balanced Salt Solution (HBSS). The cortex hemispheres were separated and the meningeal membranes removed. Cortices were transferred to a dish with HBSS. Tissue was minced with small scissors and incubated with 2ml trypsin (0.05%) and DNase (7500U, 5mg/ml Sigma) at 37^0^C for 5min before further dissociating the cell suspension using a glass pipette. 8ml medium (DMEM, D6429, Sigma, with 10% heat-inactivated FBS, F7524, Sigma) and penicillin/ streptomycin) was added to inhibit the trypsin. The suspension was then centrifuged at 1400rpm for 5mins and the supernatant removed. The cell pellet was resuspended in 6ml medium and transferred to poly-D-lysine-coated (P0899, Sigma) T25 flasks (1 brain per T25 flask) and incubated at 37°C. Media was changed the following day and then alternate days until a confluent layer of astrocytes with a layer of microglia on top were visible. After 7-10 days microglia were removed from the mixed glial culture by tapping the flask against a glass bottle until the microglia detached. Microglia were then replated on poly-D-lysine-coated 6 well plates containing coverslips and used for immunofluorescence.

### Generation of human iPSC-derived microglia

Microglia were differentiated from the human induced pluripotent stem cell (iPSC) line, KOLF2.1J^67^. The iPSCs were cultured to 70–80% confluency and enzymatically dissociated using Accutase (StemCell Technologies, 07922) at 37 °C for 10 minutes, before seeding at 10,000 cells per well of a 96-well Clear Round Bottom Ultra-Low Attachment Microplate (Corning, 7007) in mTeSR media (Stemcell Technologies, 100-0276) containing 10 uM ROCKi (Sigma-Aldrich, SCM075), 50 ng/mL BMP-4 (PeproTech), 50 ng/mL VEGF (PeproTech) and 50 ng/mL SCF (PeproTech, 120-05). Treatment with BMP4 induced mesoderm, VEGF endothelial precursers and SCF hematopoietic precursors, resulting in the formation of embryoid bodies (EBs).

From day 2 EBs were cultured in media without ROCKi and on day 4 around 30 EBs were transferred into a T25 cell culture flask containing macrophage-directed differentiation media consisting of X-vivo 15 (Lonza, BE02-060F) supplemented with 100 ng/mL M-CSF, 25 ng/mL IL-3, 2 mM GlutaMAX, 0.05 mM B-mercaptoethanol (Thermo Fisher Scientific, 31350010) and 1x Antibiotic-Antimycotic (Thermo Fisher Scientific, 15240062). After a week, a complete media change was performed, thereafter only two-thirds of the media was changed every 4-5 days. The EBs attached to the plate, were surrounded adherent stromal cells and formed cystic structures reminiscent of yolk sac morphology.

Macrophage precursors emerged in the culture supernatant after 2–3 weeks and were collected at 4–5-day intervals during weeks 3 to 5 of differentiation. Macrophage precursors were seeded onto coverslips at a density of 200,000 cells per well of a 12-well plate. To promote the maturation of macrophage precursors into a microglial phenotype, cells were cultured for 14 days in a differentiation medium composed of DMEM/F12 supplemented with 1× N2 (Thermo Fisher Scientific, 17502038), 2 mM GlutaMAX (Thermo Fisher Scientific, 35050038), 50 μM 2-mercaptoethanol, 100 ng/mL interleukin-34 (PeproTech, 200-34),10 ng/mL granulocyte-macrophage colony-stimulating factor (Thermo Fisher Scientific, 2907249), and 50 U/mL Antibiotic-Antimycotic. The cultures were replenished with fresh medium every 4–5 days. Macrophages matured into microglia after 14 days in culture.

### Immunofluorescence microscopy

Cells were grown on coverslips, washed with 1x PBS and fixed in 4% paraformaldehyde for 20 min. After washing with PBS cells were permeabilized for 5 min with 0.2% Triton X-100 and blocked 1h with 1% BSA in PBS before incubating with the indicated primary antibodies for at least 1h at RT or at 4°C overnight. Primary antibodies were detected by incubating for 1h with AlexaFluor488- or AlexaFluor568-coupled secondary antibodies (Thermo Fisher Scientific). F-actin was visualised using phalloidin coupled to either AlexaFluor568 or AlexaFluor647 (Thermo Fisher Scientific, A12380, A22287) and cell nuclei were stained with Hoechst (Thermo Fisher Scientific, 33342). Cells were mounted on slides using ProLong Antifade reagent (Thermo Fisher Scientific, P36936), imaged using the Zeiss SP880 confocal microscope with or without Airyscan and images processed using FIJI.

### Immunoprecipitation and Western blotting

Immunoprecipitations were performed from RPE WT or stable cell lines. 24h after transfection cells were lysed in 50mM Tris pH7.4, 100mM NaCl, 1% NP-40, 5mM MgCl_2_, 5mM ATP and complete protease inhibitor cocktail (Roche). After shearing with a 25-gauge needle, lysates were clarified by centrifugation at 20,000g for 15min at 4^0^C. Supernatants were precleared with protein A Sepharose followed by incubation with 5µg antibody for 1h, then protein A Sepharose for 1h. After washing the beads 3 times with lysis buffer and once with PBS, proteins were boiled in SDS loading buffer and separated by SDS-PAGE. After protein transfer on to nitrocellulose (Protran, Amersham), the membrane was blocked with 5% milk in PBS-T (0.05% Tween-20 in PBS), incubated with primary antibodies overnight at 4°C, washed with PBS-T and incubated with the corresponding HRP-conjugated secondary antibody for 1h at RT. After washing, bound antibody was detected using enhanced chemiluminescence (ECL) substrate (GE Healthcare Life Sciences) according to the manufacturer’s protocol and exposed to X-ray film (Fujifilm) or imaged using a G:BOX imaging system (Syngene).

### BioID purification and sample processing

RPE cells (3x150mm dishes at 50% confluency) or U937 cells (approximately 4x10^7^ cells) were supplemented with 50µM biotin in growth media and incubated for 24h.

Cells were lysed with RIPA lysis buffer (50mM Tris-HCl pH7.4, 150mM NaCl, 1% NP-40, 0.5% sodium deoxycholate, 1mM EDTA, 0.1% SDS and complete protease inhibitor cocktail (Roche cOmplete Mini, EDTA-free)), sonicated, centrifuged, and the supernatants mixed with high capacity streptavidin beads (Thermo Fisher Scientific) for 3h at 4^0^C. Beads were washed with RIPA buffer, TBS and 50mM ammonium bicarbonate pH 8 before incubation with 10mM DTT for 30min at 56^0^C. The solution was spiked with 10µl 550mM Iodoacetamide (Sigma BioUltra) for 20min at room temperature and the beads washed in ammonium bicarbonate before digestion overnight with 0.5µg Trypsin Gold (Promega) at 37^0^C. An additional 0.5µg trypsin was added the following day and incubated for a further 2h. After centrifugation the supernatants were collected and the beads washed twice with 150µl HPLC-grade H_2_O (Sigma, CHROMASOLV) and all supernatants combined. These were spiked with 1µl Trifluoroacetic acid and dried to a pellet in a vacuum centrifuge.

### Mass spectroscopy (MS) acquisition and data analysis

Samples were resuspended in MS solvent (3% acetonitrile, 0.1% TFA) for analysis on a Q Exactive Plus (Thermo Fisher Scientific) coupled to an RSLC3000nano UPLC (Thermo Fisher Scientific). Peptides were resolved using a 50 cm C18 PepMap EASYspray column with a gradient rising from 97% solvent A (0.1% formic acid), 3% solvent B (80% acetonitrile, 0.1% formic acid) to 40% solvent B over 40 min. Data were acquired in a top 10 data-dependent acquisition fashion with MS spectra acquired between m/z 400 and 1,500 at 70,000 fwhm. MSMS spectra were acquired at 17,500 fwhm and excluded from further fragmentation for 30 s. Raw files were processed as a single batch using the MaxQuant proteomics software package version 2.4.13.0^68^. Spectra were searched against a reviewed Uniprot homo sapiens database (downloaded 24/10/23). Cysteine carbamidomethylation was set as a fixed modification, and methionine oxidation and N-terminal acetylation were selected as variable modifications. Both peptide and protein false discovery rates (FDRs) were set to 0.01, the minimum peptide length was set at 8 amino acids, and up to two missed cleavages were tolerated.

Bioinformatics analysis was performed in the Perseus package bundled with MaxQuant^69^. Data were filtered by removing matches to the reverse database, proteins only identified with modified peptides, and common contaminants and intensity values were log_10_ transformed.

Volcano plots were generated using Perseus v. 2.0.11. LFQ intensities were filtered for reverse, potential contaminant, identified by site and transformed by log_2_(x). Proteins were filtered based on minimum of 3 valid values and missing values imputed with a width of 0.3 and downshift of 2.0. The false discovery rate was set at 0.01 and the S0 value at 2.0.

### Phagocytosis assay and image Acquisition

RAW cells (approximately 400,000 cells) were nucleofected with the Lonza Nucleofector system (SF Cell Line, Lonza). Cells were pelleted by centrifugation at 90 × g 5min and resuspended in 20 µL of Nucleofector solution supplemented with 0.4 µg of plasmid DNA. Following nucleofection, cells were incubated for 5 minutes at room temepraturebefore being transferred to pre-warmed complete culture medium and seeded onto ethanol-sterilized glass coverslips for overnight culture.

BSA-coated polysterene beads (6-8µm, Spherotech BP-60-5) were washed three times with 1x PBS and incubated overnight at 4 °C with 10 µg/mL Alexa Fluor 647-conjugated anti-BSA antibody (Thermo Fisher Scientific, A7816). After 24h, the beads were washed three times with 1xPBS and resuspended in pre-warmed culture media. To initiate phagocytosis, the cell media was replaced with the prepared bead solution (10x excess over number of cells seeded) and incubated at 37°C, 5%CO_2_ for 15min before fixing in 4% PFA and 3x washing with 1x PBS.

The cells were imaged on a spinning-disc confocal microscope (Revolution; Andor) on DMi8 (Leica) housing, equipped with a spinning-disk unit (CSU-X1; Yokogawa) using a 60x Objective 1.3NA (Leica) and 405, 488 and 561nm lasers. Image stacks were acquired as averages of 2 at a z-interval of 0.2µm with an iXon Ultra 888 EmCCD camera (Andor) set to 250 Gain and 50ms exposure.

### Image Analysis

ImageJ was used to calculate maximum Z projections of the image stacks. To quantify the accumulation of protein intensity at the phagocytic cup, masks for the three channels (bead, actin and the protein of interest) were created via the OTSU algorithm. For the actin mask the threshold was set at 2* the calculated Otsu threshold. To segment the area of the actin which forms the phagocytic cup, the bead mask and the actin mask were multiplied to select only actin structures that did contain a fluorescent bead (see Suppl. Fig. 1). The accumulation of the protein of interest was then quantified as the ratio of average protein intensity inside the actin ring (I_inside_) divided by the average protein intensity in the protein mask without the actin ring (I_outside_) minus one.

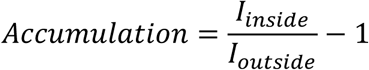

For the statistical analysis two-sided student t-tests were performed to gather the reported p values (Prism 10.1.2, Graphpad).

## REFERENCES

1. Fili N, Toseland CP. Unconventional Myosins: How Regulation Meets Function. Int J Mol Sci. 2019;21(1).

2. Hartman MA, Spudich JA. The myosin superfamily at a glance. J Cell Sci. 2012;125(Pt 7):1627–1632.

3. Masters TA, Kendrick-Jones J, Buss F. Myosins: Domain Organisation, Motor Properties, Physiological Roles and Cellular Functions. Handb Exp Pharmacol. 2017;235:77–122.

4. McIntosh BB, Ostap EM. Myosin-I molecular motors at a glance. J Cell Sci. 2016;129(14):2689–2695.

5. Giron-Perez DA, Piedra-Quintero ZL, Santos-Argumedo L. Class I myosins: Highly versatile proteins with specific functions in the immune system. J Leukoc Biol. 2019;105(5):973–981.

6. Greenberg MJ, Ostap EM. Regulation and control of myosin-I by the motor and light chain-binding domains. Trends Cell Biol. 2013;23(2):81–89.

7. Foth BJ, Goedecke MC, Soldati D. New insights into myosin evolution and classification. Proc Natl Acad Sci U S A. 2006;103(10):3681–3686.

8. McConnell RE, Tyska MJ. Leveraging the membrane - cytoskeleton interface with myosin-1. Trends Cell Biol. 2010;20(7):418–426.

9. Stoffler HE, Ruppert C, Reinhard J, Bahler M. A novel mammalian myosin I from rat with an SH3 domain localizes to Con A-inducible, F-actin-rich structures at cell-cell contacts. J Cell Biol. 1995;129(3):819–830.

10. Navines-Ferrer A, Martin M. Long-Tailed Unconventional Class I Myosins in Health and Disease. Int J Mol Sci. 2020;21(7).

11. Liu PJ, Sayeeda K, Zhuang C, Krendel M. Roles of myosin 1e and the actin cytoskeleton in kidney functions and familial kidney disease. Cytoskeleton (Hoboken). 2024;81(12):737–752.

12. Kim SV, Mehal WZ, Dong X, et al. Modulation of cell adhesion and motility in the immune system by Myo1f. Science. 2006;314(5796):136-139.

13. Salvermoser M, Pick R, Weckbach LT, et al. Myosin 1f is specifically required for neutrophil migration in 3D environments during acute inflammation. Blood. 2018;131(17):1887–1898.

14. Zhang B, Gaiteri C, Bodea LG, et al. Integrated systems approach identifies genetic nodes and networks in late-onset Alzheimer’s disease. Cell. 2013;153(3):707–720.

15. Matarin M, Salih DA, Yasvoina M, et al. A genome-wide gene-expression analysis and database in transgenic mice during development of amyloid or tau pathology. Cell Rep. 2015;10(4):633–644.

16. Gao C, Jiang J, Tan Y, Chen S. Microglia in neurodegenerative diseases: mechanism and potential therapeutic targets. Signal Transduct Target Ther. 2023;8(1):359.

17. Zhang J, Velmeshev D, Hashimoto K, et al. Neurotoxic microglia promote TDP-43 proteinopathy in progranulin deficiency. Nature. 2020;588(7838):459-465.

18. Mukherjee S, Klaus C, Pricop-Jeckstadt M, Miller JA, Struebing FL. A Microglial Signature Directing Human Aging and Neurodegeneration-Related Gene Networks. Front Neurosci. 2019;13:2.

19. Piedra-Quintero ZL, Serrano C, Villegas-Sepulveda N, et al. Myosin 1F Regulates M1-Polarization by Stimulating Intercellular Adhesion in Macrophages. Front Immunol. 2018;9:3118.

20. Cervero P, Himmel M, Kruger M, Linder S. Proteomic analysis of podosome fractions from macrophages reveals similarities to spreading initiation centres. Eur J Cell Biol. 2012;91(11-12):908–922.

21. Ouderkirk JL, Krendel M. Myosin 1e is a component of the invadosome core that contributes to regulation of invadosome dynamics. Exp Cell Res. 2014;322(2):265–276.

22. Linder S, Cervero P, Eddy R, Condeelis J. Mechanisms and roles of podosomes and invadopodia. Nat Rev Mol Cell Biol. 2023;24(2):86–106.

23. Alonso F, Spuul P, Daubon T, Kramer I, Genot E. Variations on the theme of podosomes: A matter of context. Biochim Biophys Acta Mol Cell Res. 2019;1866(4):545–553.

24. Boateng LR, Huttenlocher A. Spatiotemporal regulation of Src and its substrates at invadosomes. Eur J Cell Biol. 2012;91(11-12):878–888.

25. Linder S, Nelson D, Weiss M, Aepfelbacher M. Wiskott-Aldrich syndrome protein regulates podosomes in primary human macrophages. Proc Natl Acad Sci U S A. 1999;96(17):9648–9653.

26. Burgstaller G, Gimona M. Actin cytoskeleton remodelling via local inhibition of contractility at discrete microdomains. J Cell Sci. 2004;117(Pt 2):223–231.

27. Moreau V, Tatin F, Varon C, Genot E. Actin can reorganize into podosomes in aortic endothelial cells, a process controlled by Cdc42 and RhoA. Mol Cell Biol. 2003;23(19):6809–6822.

28. Zhang Y, Cao F, Zhou Y, et al. Tail domains of myosin-1e regulate phosphatidylinositol signaling and F-actin polymerization at the ventral layer of podosomes. Mol Biol Cell. 2019;30(5):622–635.

29. Herron JC, Hu S, Watanabe T, et al. Actin nano-architecture of phagocytic podosomes. Nat Commun. 2022;13(1):4363.

30. Linder S, Barcelona B. Get a grip: Podosomes as potential players in phagocytosis. Eur J Cell Biol. 2023;102(4):151356.

31. Vorselen D, Barger SR, Wang Y, et al. Phagocytic ’teeth’ and myosin-II ’jaw’ power target constriction during phagocytosis. Elife. 2021;10.

32. Barger SR, Vorselen D, Gauthier NC, Theriot JA, Krendel M. F-actin organization and target constriction during primary macrophage phagocytosis is balanced by competing activity of myosin-I and myosin-II. Mol Biol Cell. 2022;33(14):br24.

33. Barger SR, Reilly NS, Shutova MS, et al. Membrane-cytoskeletal crosstalk mediated by myosin-I regulates adhesion turnover during phagocytosis. Nat Commun. 2019;10(1):1249.

34. Patino-Lopez G, Aravind L, Dong X, Kruhlak MJ, Ostap EM, Shaw S. Myosin 1G is an abundant class I myosin in lymphocytes whose localization at the plasma membrane depends on its ancient divergent pleckstrin homology (PH) domain (Myo1PH). J Biol Chem. 2010;285(12):8675–8686.

35. Brown MT, Andrade J, Radhakrishna H, Donaldson JG, Cooper JA, Randazzo PA. ASAP1, a phospholipid-dependent arf GTPase-activating protein that associates with and is phosphorylated by Src. Mol Cell Biol. 1998;18(12):7038–7051.

36. Luo R, Reed CE, Sload JA, Wordeman L, Randazzo PA, Chen PW. Arf GAPs and molecular motors. Small GTPases. 2019;10(3):196–209.

37. Take H, Watanabe S, Takeda K, Yu ZX, Iwata N, Kajigaya S. Cloning and characterization of a novel adaptor protein, CIN85, that interacts with c-Cbl. Biochem Biophys Res Commun. 2000;268(2):321–328.

38. Dikic I. CIN85/CMS family of adaptor molecules. FEBS Lett. 2002;529(1):110–115.

39. Kirsch KH, Georgescu MM, Ishimaru S, Hanafusa H. CMS: an adapter molecule involved in cytoskeletal rearrangements. Proc Natl Acad Sci U S A. 1999;96(11):6211–6216.

40. Deckert M, Rottapel R. The adapter 3BP2: how it plugs into leukocyte signaling. Adv Exp Med Biol. 2006;584:107–114.

41. Ueki Y, Tiziani V, Santanna C, et al. Mutations in the gene encoding c-Abl-binding protein SH3BP2 cause cherubism. Nat Genet. 2001;28(2):125–126.

42. Navines-Ferrer A, Ainsua-Enrich E, Serrano-Candelas E, Sayos J, Martin M. Myo1f, an Unconventional Long-Tailed Myosin, Is a New Partner for the Adaptor 3BP2 Involved in Mast Cell Migration. Front Immunol. 2019;10:1058.

43. Derry JM, Ochs HD, Francke U. Isolation of a novel gene mutated in Wiskott-Aldrich syndrome. Cell. 1994;79(5):following 922.

44. Lee PP, Lobato-Marquez D, Pramanik N, et al. Wiskott-Aldrich syndrome protein regulates autophagy and inflammasome activity in innate immune cells. Nat Commun. 2017;8(1):1576.

45. Aspenstrom P. Formin-binding proteins: modulators of formin-dependent actin polymerization. Biochim Biophys Acta. 2010;1803(2):174–182.

46. Katoh M, Katoh M. Identification and characterization of the human FMN1 gene in silico. Int J Mol Med. 2004;14(1):121–126.

47. Wu Z, Blessing NA, Simske JS, Bruggeman LA. Fyn-binding protein ADAP supports actin organization in podocytes. Physiol Rep. 2017;5(23).

48. Coppolino MG, Krause M, Hagendorff P, et al. Evidence for a molecular complex consisting of Fyb/SLAP, SLP-76, Nck, VASP and WASP that links the actin cytoskeleton to Fcgamma receptor signalling during phagocytosis. J Cell Sci. 2001;114(Pt 23):4307–4318.

49. Takemitsu M, Ishiura S, Koga R, et al. Dystrophin-related protein in the fetal and denervated skeletal muscles of normal and mdx mice. Biochem Biophys Res Commun. 1991;180(3):1179–1186.

50. Winder SJ, Hemmings L, Maciver SK, et al. Utrophin actin binding domain: analysis of actin binding and cellular targeting. J Cell Sci. 1995;108 (Pt 1):63–71.

51. Citterio L, Azzani T, Duga S, Bianchi G. Genomic organization of the human gamma adducin gene. Biochem Biophys Res Commun. 1999;266(1):110–114.

52. Georges-Labouesse E, Mark M, Messaddeq N, Gansmuller A. Essential role of alpha 6 integrins in cortical and retinal lamination. Curr Biol. 1998;8(17):983–986.

53. Newman PJ, Berndt MC, Gorski J, et al. PECAM-1 (CD31) cloning and relation to adhesion molecules of the immunoglobulin gene superfamily. Science. 1990;247(4947):1219–1222.

54. Feng YM, Chen XH, Zhang X. Roles of PECAM-1 in cell function and disease progression. Eur Rev Med Pharmacol Sci. 2016;20(19):4082–4088.

55. Kuninty PR, Bansal R, De Geus SWL, et al. ITGA5 inhibition in pancreatic stellate cells attenuates desmoplasia and potentiates efficacy of chemotherapy in pancreatic cancer. Sci Adv. 2019;5(9):eaax2770.

56. Lai-Cheong JE, Parsons M, McGrath JA. The role of kindlins in cell biology and relevance to human disease. Int J Biochem Cell Biol. 2010;42(5):595–603.

57. Farrugia AJ, Calvo F. The Borg family of Cdc42 effector proteins Cdc42EP1-5. Biochem Soc Trans. 2016;44(6):1709–1716.

58. Bharti S, Inoue H, Bharti K, et al. Src-dependent phosphorylation of ASAP1 regulates podosomes. Mol Cell Biol. 2007;27(23):8271–8283.

59. Shiba Y, Randazzo PA. GEFH1 binds ASAP1 and regulates podosome formation. Biochem Biophys Res Commun. 2011;406(4):574–579.

60. Maxeiner S, Shi N, Schalla C, et al. Crucial role for the LSP1-myosin1e bimolecular complex in the regulation of Fcgamma receptor-driven phagocytosis. Mol Biol Cell. 2015;26(9):1652–1664.

61. Vadillo E, Chanez-Paredes S, Vargas-Robles H, et al. Intermittent rolling is a defect of the extravasation cascade caused by Myosin1e-deficiency in neutrophils. Proc Natl Acad Sci U S A. 2019;116(52):26752–26758.

62. McCauley ME, O’Rourke JG, Yanez A, et al. C9orf72 in myeloid cells suppresses STING-induced inflammation. Nature. 2020;585(7823):96–101.

63. Vahsen BF, Nalluru S, Morgan GR, et al. C9orf72-ALS human iPSC microglia are pro-inflammatory and toxic to co-cultured motor neurons via MMP9. Nat Commun. 2023;14(1):5898.

64. Hollingworth P, Harold D, Sims R, et al. Common variants at ABCA7, MS4A6A/MS4A4E, EPHA1, CD33 and CD2AP are associated with Alzheimer’s disease. Nat Genet. 2011;43(5):429–435.

65. Naj AC, Jun G, Beecham GW, et al. Common variants at MS4A4/MS4A6E, CD2AP, CD33 and EPHA1 are associated with late-onset Alzheimer’s disease. Nat Genet. 2011;43(5):436–441.

66. Havrylov S, Redowicz MJ, Buchman VL. Emerging roles of Ruk/CIN85 in vesicle-mediated transport, adhesion, migration and malignancy. Traffic. 2010;11(6):721–731.

67. Pantazis CB, Yang A, Lara E, et al. A reference human induced pluripotent stem cell line for large-scale collaborative studies. Cell Stem Cell. 2022;29(12):1685–1702 e1622.

68. Cox J, Mann M. MaxQuant enables high peptide identification rates, individualized p.p.b.-range mass accuracies and proteome-wide protein quantification. Nat Biotechnol. 2008;26(12):1367–1372.

69. Tyanova S, Temu T, Sinitcyn P, et al. The Perseus computational platform for comprehensive analysis of (prote)omics data. Nat Methods. 2016;13(9):731–740.

