## supplementary figure 1-3 and table 1 and 2 for "MYO1F interactome reveals the SH3-domain linked CASS complex at podosomes and the phagocytic cup"

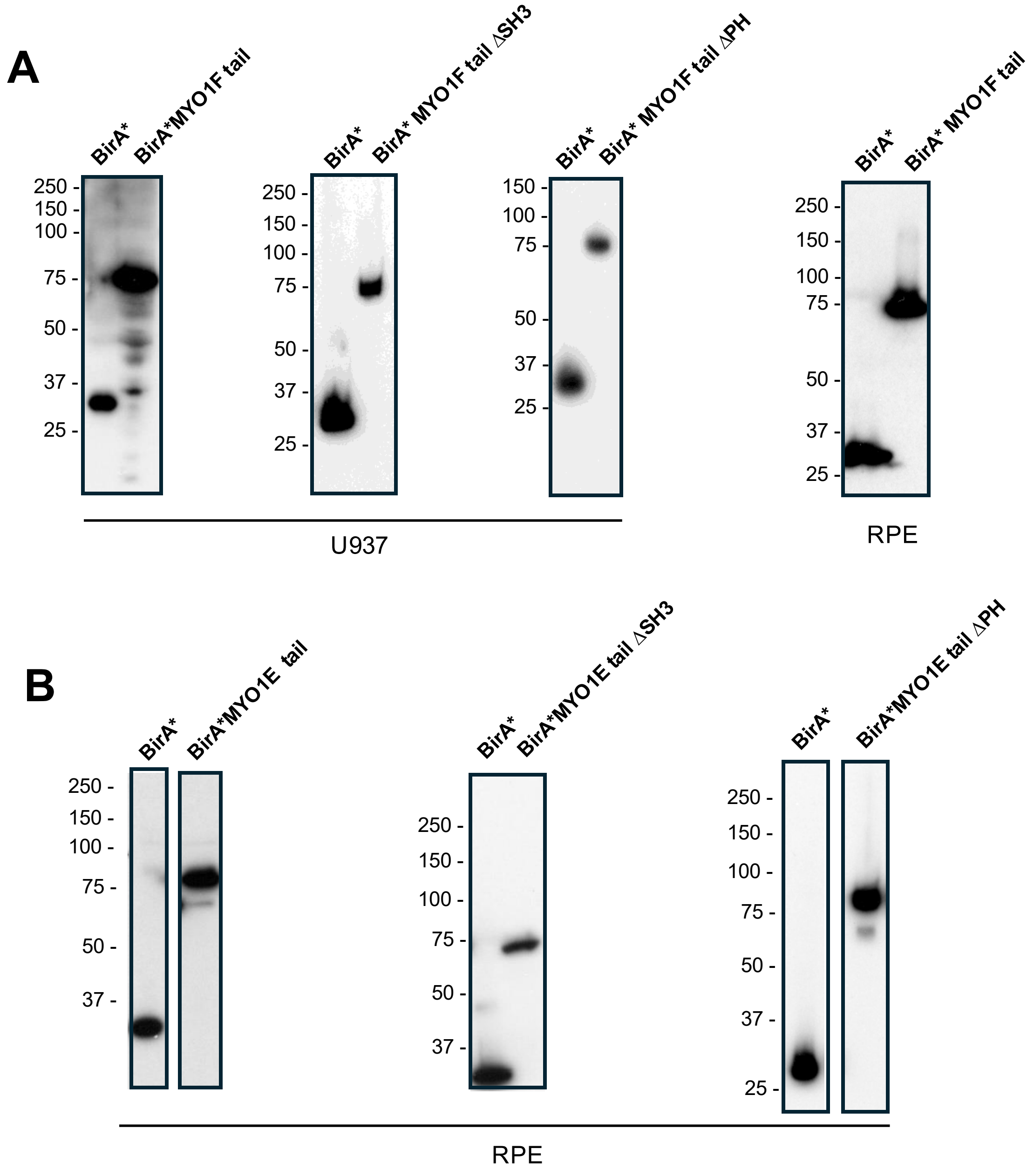
A
BirA*
BirA*MYO1F tail
BirA*
BirA* MYO1F tail ΔSH3
BirA*
BirA* MYO1F tail ΔPH
BirA*
BirA* MYO1F tail
250 -
150 -
100 -
75 -
50 -
37 -
25 -
U937
RPE
B
BirA*
BirA*MYO1E tail
BirA*
BirA*MYO1E tail ΔSH3
BirA*
BirA*MYO1E tail ΔPH
RPE

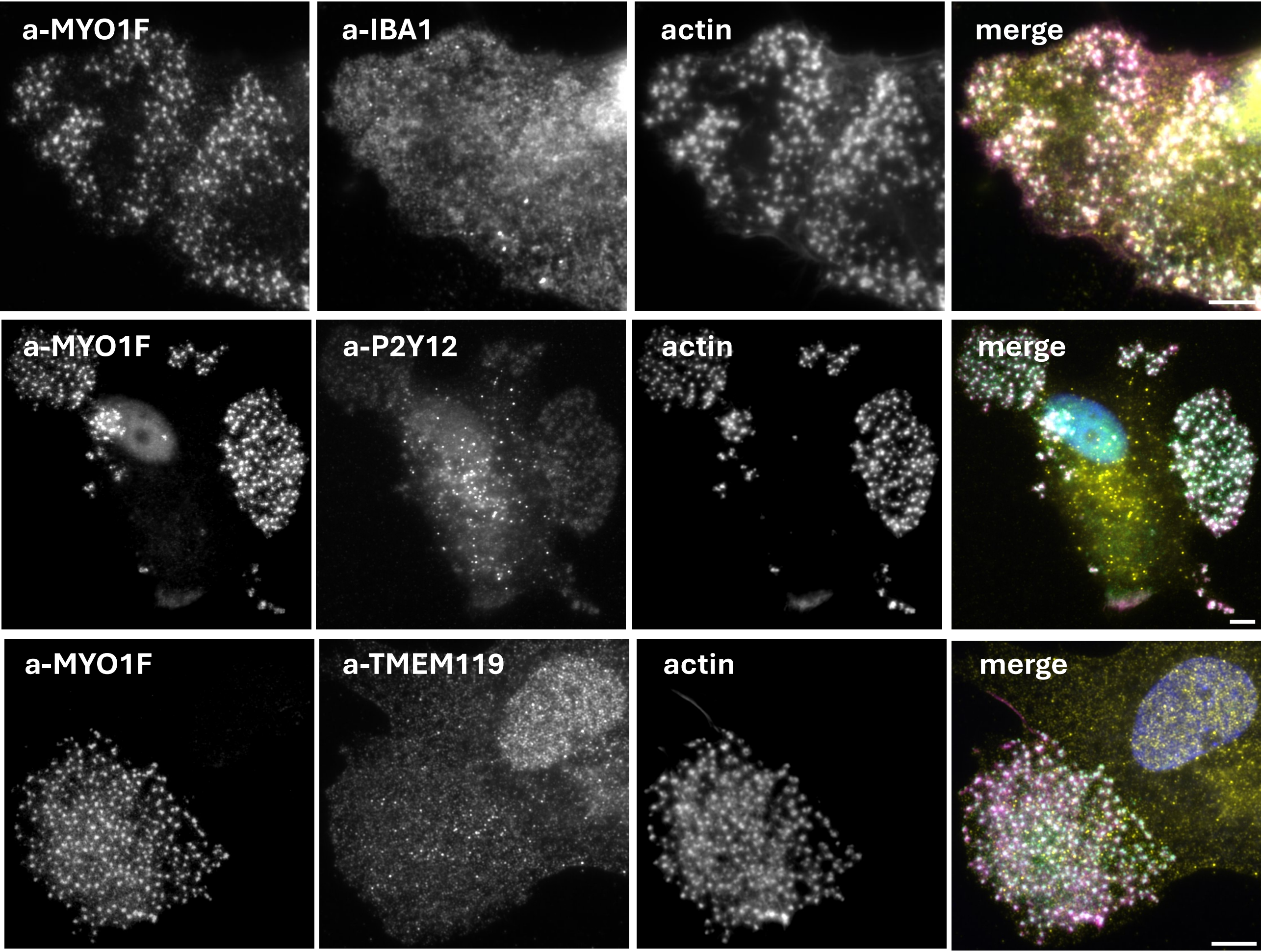
a-MYO1F
a-IBA1
actin
merge
a-MYO1F
a-P2Y12
actin
merge
a-MYO1F
a-TMEM119
actin
merge

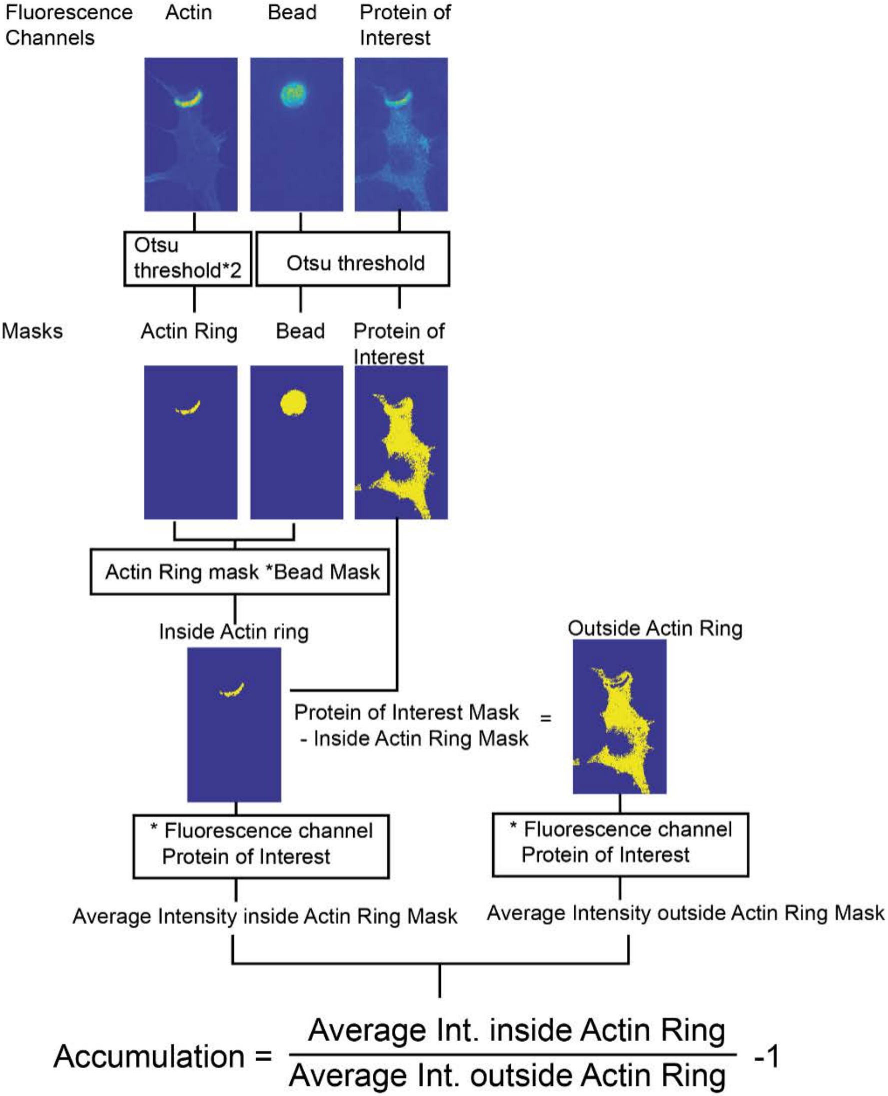
Fluorescence Channels
Actin
Bead
Protein of Interest
Otsu threshold*2
Otsu threshold
Masks
Actin Ring
Bead
Protein of Interest
Actin Ring mask *Bead Mask
Inside Actin ring
Outside Actin Ring
Protein of Interest Mask - Inside Actin Ring Mask =
* Fluorescence channel Protein of Interest
* Fluorescence channel Protein of Interest
Average Intensity inside Actin Ring Mask
Average Intensity outside Actin Ring Mask
Accumulation = Average Int. inside Actin Ring / Average Int. outside Actin Ring -1

| | Myo1F-control | | | MYO1F ΔSH3-control | | | MYO1F ΔPH-control | |
|---|---|---|---|---|---|---|---|---|
| Gene name | Difference | -Log p value | Gene nam | Difference | -Log p value | Gene nam | Difference | -Log p value |
| MYO1F | 95182724 | 168144603 | MYO1F | 8428194682 | 117680394 | MYO1F | 1019820277 | 1271999997 |
| PHACTR4 | 7012141546 | 975843702 | ACTB | 742597035 | 174151808 | PHACTR4 | 6067904154 | 6033972222 |
| PLEKHO2 | 6962905066 | 15689194 | SPTBN1 | 7032675561 | 356594429 | PRRC2A | 5955808367 | 9536057126 |
| ERBB2IP | 6273739588 | 988698146 | ACTC1;ACT | 6901574952 | 12717178 | NAP1L4 | 5778435616 | 3032724566 |
| ASAP1 | 6225703512 | 106559299 | SPTAN1 | 6899079209 | 32190874 | CSDE1 | 5744787807 | 3143148217 |
| DOCK8 | 6075487273 | 942410928 | IQGAP1 | 6435603142 | 247675035 | ASAP1 | 5456758772 | 7057171045 |
| ZDHHC5 | 582905129 | 106886317 | WDR1 | 6415785154 | 466748051 | NAP1L1 | 5443313962 | 270515481 |
| SH3BP2 | 5745374316 | 988560422 | ACTN4 | 6260526566 | 156191274 | YTHDF2 | 5434397697 | 6515966115 |
| ESYT1 | 5594581559 | 663350921 | ERBB2IP | 6181767555 | 826503687 | NUFIP2 | 5286873318 | 3266921066 |
| DLG1 | 5431517601 | 100963292 | FAM129B | 6119864509 | 992684419 | XRN1 | 5281040101 | 3033719559 |
| NUMB | 5293050675 | 102169884 | ACTN1 | 593949268 | 202454008 | SH3BP2 | 5203657423 | 6803604526 |
| MAP4K4 | 5270052637 | 863014305 | RASA3 | 5653010959 | 882255415 | FAM120A | 5183748881 | 6694038671 |
| UTRN | 5252506211 | 626102963 | CORO1C | 5602709044 | 282071283 | IPO7 | 5157002631 | 9624697616 |
| FAM129B | 5168643361 | 10130213 | AIF1 | 5594447227 | 314061182 | PLEKHO2 | 5120529311 | 9895447269 |
| LAT2 | 5149946939 | 562151902 | TMOD3 | 5574904805 | 239615617 | PCM1 | 5086153621 | 2762660038 |
| ESYT2 | 5121908052 | 689238609 | LIMA1 | 5568374952 | 315168531 | PRRC2C | 5067368144 | 4919736614 |
| SH3KBP1 | 5108538764 | 464668458 | ZDHHC5 | 5221820786 | 858801889 | UBAP2 | 5029208955 | 4292376364 |
| PPFIBP2 | 509948072 | 118307791 | DLG1 | 5170201302 | 732291072 | UBAP2L | 497335216 | 2568175748 |
| SLC4A7 | 5012280192 | 618087535 | MYO1G | 5119916235 | 239557266 | ZDHHC5 | 4960267339 | 8268946067 |
| RASA3 | 4854565257 | 734299371 | PVRL2 | 5116679419 | 650143911 | TDRD3 | 4903487932 | 7366700256 |
| CD99 | 4823954128 | 385092752 | TWF2 | 511103344 | 204936518 | LCP2 | 4893806094 | 3874764359 |
| STEAP3 | 4815965652 | 489951259 | CAPZA2 | 5098364785 | 176667323 | DVL1;DVL1 | 482086554 | 410786493 |
| MARK3 | 4786682174 | 105367732 | ITGA6 | 5052712758 | 307295981 | ATXN2L | 4728120213 | 2133696033 |
| SLC30A1 | 4651078769 | 636812096 | PHACTR4 | 5026692708 | 505449829 | UBE2O | 4726200104 | 3427872332 |
| SIRPA;SIRPB1 | 4625443595 | 604666009 | SLC38A1 | 4995862053 | 760572407 | LARP1 | 4702118374 | 2019588885 |
| AHCYL1 | 4611974762 | 652200376 | ACTG1 | 4987374669 | 215230104 | ANKRD17 | 4701183228 | 4509791218 |
| GPR124 | 4602253505 | 939975987 | PVRL1 | 4981006622 | 827411333 | STRAP | 465124598 | 2749786059 |
| CD2AP | 4560138566 | 405441569 | ESYT1 | 4980293546 | 391474634 | MAP7D3 | 4646885009 | 6547724391 |
| WAS | 4540426254 | 369475073 | CFL1 | 4954517819 | 218819866 | YTHDF3 | 4630905469 | 90620197 |
| SLC7A5 | 4518964677 | 386661758 | DOCK8 | 4881989615 | 570844164 | WAS | 4598633448 | 2328625545 |
| EFR3A | 4417370115 | 778876993 | CD44 | 4856786138 | 339664006 | MAP4K5 | 4585683959 | 6814593005 |
| BASP1 | 4371285484 | 881360967 | MAP4K4 | 4831797327 | 567551244 | EIF3L | 4572031475 | 3745500827 |
| ADD3 | 4357732773 | 101055435 | RELL1 | 4827170554 | 72866191 | UPF1 | 4568852016 | 5189616728 |
| MARK2 | 4287561553 | 722297426 | TWF1 | 4809035437 | 321832622 | MAP4K4 | 4473332133 | 5299594546 |
| NF2 | 4284461339 | 87408155 | LAT2 | 4752916109 | 331138557 | SND1 | 4461706025 | 2209418085 |
| GOLGA8R | 4270157632 | 244595646 | NDRG1 | 4659833817 | 257502836 | LSM12 | 4461231822 | 2661917436 |
| CD44 | 4255859103 | 326276127 | EFR3A | 4650655701 | 647964551 | ATXN2 | 4454963775 | 6653720494 |
| CDCA3 | 4188340142 | 127359067 | MARK3 | 4571919487 | 787067869 | ZC3HAV1 | 4438353584 | 2832073806 |
| ITGA6 | 4151344935 | 37821336 | BASP1 | 4502928779 | 6758181 | EIF4B | 4433714276 | 1625043399 |
| FNBP1 | 4115563393 | 684891355 | CD99 | 4467542194 | 218331487 | ZCCHC6 | 4433073725 | 7519169467 |
| SNAP29 | 4096730823 | 456001271 | MCEMP1 | 4376711709 | 487225182 | ERBB2IP | 4413307599 | 6312776809 |
| CDC37 | 4078293664 | 41178837 | CAPZB | 436408488 | 199113182 | SERBP1 | 4370198976 | 2739691139 |
| FYB | 4074363618 | 469032301 | ESYT2 | 4361855689 | 391353366 | GEMIN5 | 4304373469 | 1573241832 |
| KANK2 | 4061893826 | 337219161 | NF2 | 4347975413 | 675699548 | NACA | 4291952587 | 215665394 |
| SH3GL1 | 4059275718 | 511733855 | CORO2B | 4305422465 | 402553308 | EIF4G1 | 4277588027 | 173763578 |
| SIGLEC6 | 4040959086 | 390058756 | | 4299947012 | 236199838 | SH3KBP1 | 4230715252 | 2306475351 |
| PECAM1 | 4016202382 | 904549691 | LRRC25 | 4289059821 | 79001981 | GIGYF2 | 4218240556 | 2363054886 |
| MARCKS | 401343936 | 648235954 | SIRPA;SIRP | 4281774339 | 546039799 | EIF3B | 4169646172 | 2445693078 |
| NDRG1 | 3972479412 | 338504295 | FCER1G | 4266245161 | 724349206 | EIF2A | 4121648925 | 1933837202 |
| SNAP23 | 3964447476 | 600629621 | ACTBL2 | 4255010832 | 199635087 | DVL3 | 412057209 | 2577502375 |
| SIGLEC5;SIGL | 3948899042 | 636086649 | UTRN | 4253845487 | 329650352 | CD2AP | 4101004464 | 2202649503 |
| EPB41 | 3887228012 | 862198443 | KANK2 | 4248226211 | 221033014 | SIGLEC6 | 4098211334 | 2909531179 |
| PACSIN2 | 3880630312 | 48499788 | AHCYL1 | 4123611813 | 507406375 | EIF3E | 4092319988 | 3107535151 |
| REPS1 | 3868869918 | 420934043 | NUMB | 4099826404 | 72897502 | TCHP | 4045064154 | 2506179143 |
| MAP4K5 | 3846785999 | 509360939 | SNAP23 | 4070356187 | 436070824 | ZC3H15 | 4001512437 | 2153974667 |
| LIMD1 | 3837959471 | 406536963 | CORO2A | 406830883 | 450469855 | ESYT1 | 3985702151 | 3038269637 |
| RELL1 | 3765082223 | 377196778 | ARPC3 | 4017595473 | 177190253 | EIF3G | 3957202548 | 247373707 |
| SLC3A2 | 3711643083 | 322450227 | CAPZA1 | 3886984235 | 227163218 | EIF4G2 | 3950390952 | 3119970651 |
| SNX9 | 3676011585 | 900117338 | PECAM1 | 387948236 | 74745324 | ZYX | 3906148683 | 1386313064 |
| WDR44 | 3662206514 | 648357593 | ARPC2 | 385840802 | 165102913 | PRAM1 | 3839349792 | 1936832389 |
| HMHA1 | 3645019531 | 608943123 | ARPC5 | 3854095913 | 147368777 | BRAP | 38252093 | 2083325401 |
| PRAM1 | 3623187428 | 275510882 | SLC4A7 | 3851324445 | 296056121 | DHX29 | 3808204197 | 4347970479 |
| TBC1D10B | 3615252313 | 819626545 | CPNE8 | 3825607754 | 732963528 | CRKL | 3801685424 | 1310209099 |
| JAG1 | 3610378175 | 108709685 | TMOD2 | 381439209 | 378159365 | G3BP1 | 3773753802 | 3591669318 |
| ZC3HAV1 | 3604825065 | 340927872 | FRMD3 | 3770354135 | 688620561 | EML4 | 3725784756 | 4907746251 |
| PVRL2 | 3597171693 | 399831422 | SLC30A1 | 3741694995 | 395725939 | DDX6 | 3705362229 | 2748815486 |
| GAB1 | 3516349838 | 990392879 | GPR124 | 3701374599 | 70445403 | LIMD1 | 3703510148 | 2867609603 |
| PEAK1 | 3449781236 | 592557196 | CD33 | 3675500643 | 642018416 | PACSIN2 | 3691130139 | 7073935893 |
| SBF1 | 3393878982 | 592790421 | ARPC1B | 3663477444 | 169067592 | RGL2 | 368412095 | 4921216901 |
| PPFIBP1 | 3316645214 | 822755302 | SIGLEC6 | 3655917531 | 252893543 | YTHDF1 | 367861775 | 527581445 |
| WIPF1 | 3300072625 | 429054494 | SNAP29 | 3610914503 | 266295696 | UTRN | 3670275007 | 276635346 |
| FMNL1 | 3296294621 | 38779722 | RAB11A;R/ | 360799635 | 478449941 | MAPRE2 | 3663354102 | 1620859026 |
| SEMA4D | 3291478657 | 620561877 | RASA2 | 3562689418 | 650999603 | RALBP1 | 3661553701 | 2975226306 |
| PVRL1 | 3284757296 | 346835409 | ITGA5 | 3555626006 | 244602281 | SLC7A5 | 3647180149 | 222942397 |
| SLC39A10 | 3280790011 | 2999983 | MYL6 | 3512391635 | 175432745 | R3HDM1 | 3638808023 | 6143866855 |

| | | |
|---|---|---|
| ADD1 | 3280127298 | 677371981 |
| EPB41L3 | 3266217958 | 841514041 |
| NCKAP1L | 3194268272 | 432452043 |
| HLA-A | 3190551985 | 317640381 |
| EHBP1 | 3183291163 | 579071252 |
| EPS15L1 | 3168661663 | 194073165 |
| ARHGAP30 | 316019635 | 464183218 |
| DENND3 | 3159568968 | 326776469 |
| STK10 | 3152975491 | 424745713 |
| RASAL3 | 3115138553 | 385256783 |
| SPTBN1 | 3108141082 | 191339545 |
| CYFIP1 | 3101106553 | 36787815 |
| ITGA4 | 3087145533 | 31439539 |
| SASH3 | 3086857342 | 417809068 |
| MTMR1 | 3079359781 | 482968537 |
| ITGB1 | 3069386618 | 259903222 |
| FLVCR1 | 3056066967 | 576050315 |
| BTK | 3034178507 | 292178326 |
| SEMA4C | 2993795713 | 37588784 |
| PSD4 | 2982404709 | 368240562 |
| CD300LF | 2964966093 | 362798262 |
| ELMO1 | 2957240968 | 401288549 |
| YWHAG | 2941692579 | 364202963 |
| EVI2B | 2916008904 | 296011865 |
| ATP2B4 | 291555995 | 364126701 |
| ITGA5 | 2910311154 | 280796257 |
| SLC38A1 | 2903448468 | 255296785 |
| ATP8B4 | 2884765807 | 305900657 |
| PTPRA | 2875757217 | 297592058 |
| CLDND1 | 2863726752 | 289669783 |
| SIGLEC12 | 2837178821 | 26044603 |
| ROCK1 | 2811687969 | 366945511 |
| RALBP1 | 2789056142 | 241575889 |
| MPP7 | 2768266224 | 362544107 |
| SLC16A3 | 27188995 | 375885708 |
| FCER1G | 2698544139 | 257679986 |
| BIN2 | 2683682987 | 333136569 |
| LAIR1 | 2659116654 | 244812394 |
| SLC19A1 | 2619558198 | 29088005 |
| LRCH1 | 2575132143 | 370375573 |
| YWHAE | 2570482844 | 26278919 |
| LRRC25 | 2558143479 | 277228604 |
| DEPDC1B | 2556379863 | 379318196 |
| RAB11A;RAB1 | 2525578862 | 242436287 |
| SDK1 | 2524629502 | 419783686 |
| PPFIA1 | 2505487942 | 485069974 |
| RFTN1 | 2454390571 | 275338566 |
| EFNB1 | 238419501 | 313120041 |
| CPNE8 | 2357340631 | 312577087 |
| PTPN22 | 2328005609 | 325391206 |
| PTPRC | 2301578522 | 412258081 |
| ROCK2 | 2277737527 | 570124405 |
| GOLGA8N;GC | 2265367599 | 332507854 |
| AHNAK | 2263286727 | 731422041 |

| | | |
|---|---|---|
| JAG1 | 3488159089 | 744020171 |
| SLC9A3R1 | 3484202521 | 214398136 |
| HLA-A | 3482082276 | 221409596 |
| SLC39A10 | 3463603973 | 478221482 |
| MARK2 | 3456160999 | 532901817 |
| PPFIBP2 | 3432524136 | 6033968 |
| ADD3 | 3402330399 | 889564672 |
| EPB41L3 | 3345421882 | 581564648 |
| MTMR1 | 3335140001 | 453507398 |
| EPB41 | 3271905263 | 505143372 |
| TBC1D10B | 3236129897 | 586097964 |
| STEAP3 | 3229681651 | 196023257 |
| LAIR1 | 3222387223 | 267812706 |
| PPFIBP1 | 3174585887 | 532377484 |
| SLC16A3 | 3156701815 | 357337539 |
| PAG1 | 3148984727 | 515222685 |
| MARCKS | 3146426383 | 339909208 |
| CLDND1 | 3132203238 | 580588871 |
| PLEKHO2 | 3120777902 | 648732705 |
| KDELR1 | 3104742822 | 345899562 |
| ITGA4 | 3034102485 | 295301516 |
| SNX9 | 3030323846 | 602032154 |
| KCNN4 | 3007288706 | 60774395 |
| SIGLEC5;SI | 2982292266 | 392250054 |
| RAP2C;RAI | 258924398 | 760024805 |

| | | |
|---|---|---|
| EIF4E2 | 3620175952 | 3681186362 |
| SKA3 | 3619620369 | 2475362714 |
| FAM129B | 3607855842 | 6673172311 |
| SYNJ2 | 3605391139 | 4902256892 |
| SLC38A1 | 3594672248 | 5791299702 |
| LUZP1 | 3589303652 | 2075510951 |
| HAUS6 | 358026618 | 2981383721 |
| PGAM5 | 3571344557 | 1579561033 |
| FYB | 3569508144 | 3021807027 |
| ANKRD28 | 3565747806 | 3440285877 |
| EIF3K | 356107957 | 2432940732 |
| EIF2S2 | 3551051458 | 3448062097 |
| EIF3I | 354502687 | 2167399533 |
| WIPF1 | 3535119647 | 4166938172 |
| CD44 | 3534196263 | 2345153095 |
| EIF3C;EIF3( | 3519196056 | 2366940893 |
| EPS15L1 | 351861436 | 1426842954 |
| AHCYL1 | 3506852195 | 4295809406 |
| TOP3B | 3492062932 | 9089498528 |
| DVL2 | 3491349039 | 168815097 |
| ESYT2 | 3490875108 | 3030033135 |
| LRRC25 | 3490565482 | 6802866166 |
| MAPRE1 | 3476990473 | 1855878083 |
| LAT2 | 3468278022 | 2259350182 |
| SH3GL1 | 3449184191 | 2919578818 |
| MAP4 | 3439296132 | 1484138042 |
| PPFIBP2 | 3427594911 | 6324219474 |
| HAUS4 | 339918618 | 5418854057 |
| REPS1 | 3379128274 | 2486586003 |
| CBL | 3377821014 | 2841783444 |
| HDLBP | 3364061129 | 3481702837 |
| DOCK8 | 3363551912 | 3962184408 |
| DLG1 | 3354281425 | 4854770827 |
| EIF4G3 | 3345483462 | 3104901165 |
| SNAP29 | 3337913786 | 2414934367 |
| HAUS5 | 3330346698 | 4187758815 |
| HAUS7 | 3326657613 | 3766000315 |
| EIF4E | 3321934064 | 2128972537 |
| ASCC3 | 3321101915 | 3072057509 |
| STAU2 | 3309928712 | 5541632271 |
| EVI2B | 3308066323 | 5442731483 |
| CDC37 | 3278086526 | 1933541633 |
| SKAP2 | 3276232311 | 3382946827 |
| STEAP3 | 3244706472 | 1995918714 |
| USP10 | 3234869866 | 2990742997 |
| PIK3AP1 | 3232020151 | 7181624777 |
| EIF3A | 3226962907 | 2446170213 |
| HAUS3 | 3210023244 | 4241184338 |
| DRG1 | 3206087703 | 2283428318 |
| EIF3M | 3201646941 | 2644404433 |
| KIAA0430 | 3197212401 | 6535448846 |
| BTF3 | 3185535794 | 2393309453 |
| LSM14A | 3184643064 | 5049727038 |
| SIGLEC5;SI | 3177813303 | 4215637818 |
| EIF5B | 3165723256 | 2180713132 |
| FMR1 | 3128939129 | 6650561636 |
| PVRL1 | 3122046153 | 5707515015 |
| CNOT1 | 3098198437 | 4444821596 |
| DENND3 | 3073627018 | 2272874386 |
| ELMO1 | 3071322986 | 4175728803 |
| PLK1 | 3058236576 | 4662826039 |
| CYFIP1 | 3057664145 | 2790005084 |
| EIF3F | 3050006866 | 2024983137 |
| DDX20 | 302129146 | 3229626785 |
| PPP6R1 | 3020204453 | 2785553546 |
| TRIM25 | 3012209938 | 2683582357 |
| ARHGAP3( | 2983133997 | 5805190699 |
| OFD1 | 2965091342 | 4411568656 |
| FARSA | 2910114334 | 2272072759 |
| LSG1 | 288988068 | 4849450929 |
| AP2B1 | 286989044 | 3605355422 |
| DHX57 | 2865129607 | 6167119384 |
| EFR3A | 2851531937 | 4032724407 |
| MTDH | 2824411892 | 2473624255 |
| KIF14 | 2809056827 | 423205881 |
| IFT74 | 280023157 | 2830470232 |
| RASA3 | 2782106036 | 4137401552 |
| HAUS2 | 2769178073 | 2937246888 |

| | | |
|---|---|---|
| SKA1 | 2759249051 | 388541551 |
| RELL1 | 2741166296 | 2590848144 |
| CCDC124 | 2723582177 | 5258780568 |
| SIRPA;SIRP | 2717915671 | 3374306963 |
| PVRL2 | 2710508256 | 3442196001 |
| TBC1D10B | 2666385151 | 4939375466 |
| CD300LF | 2651349658 | 3165757087 |
| LARP4B | 2601059051 | 3374336189 |
| C2CD5 | 2598056248 | 4099392474 |
| LTV1 | 2595390501 | 5262461972 |
| MARK3 | 2593221392 | 4788384991 |
| CLASP1 | 257395926 | 4585358885 |
| TNRC6B | 2508904048 | 3051191802 |
| HMHA1 | 2488037427 | 366390625 |
| AGTPBP1 | 2463837805 | 4160092793 |
| MARK2 | 245943846 | 3779332731 |
| PIK3C2B | 2420155389 | 5298495707 |
| CNOT10 | 2386745407 | 4010731792 |
| ABI1 | 2372036843 | 4034493856 |

| | Myo1E-control | | | Myo1E ΔSH3-control | | | Myo1E ΔPH-control | |
|---|---|---|---|---|---|---|---|---|
| Gene name | Difference | -Log p value | Gene name | Difference | -Log p value | Gene name | Difference | -Log p value |
| MYO1E | 1103660107 | 672392594 | MYO1E | 8464753787 | 578952122 | MYO1E | 109428091 | 684307014 |
| ITGA2 | 7422002157 | 49785258 | FAM171A1 | 7585182826 | 479838085 | ITGAV | 7232983907 | 439254353 |
| ITGA5 | 7402478536 | 669748361 | VANGL1 | 6585095406 | 600526172 | ITGA5 | 6468493144 | 681453066 |
| FAM171A1 | 6952454885 | 455309233 | RBMX;RBMXI | 6394816717 | 68250015 | ITGA2 | 6287709554 | 447542148 |
| ITGAV | 6851957321 | 373375042 | ITGAV | 6376674652 | 399885501 | FAM171A1 | 6266412099 | 422078246 |
| SNAP23 | 654679203 | 66353484 | ITGA2 | 5494361877 | 413183249 | ITGB5 | 5914620399 | 402908028 |
| NOTCH2 | 6417052587 | 603206858 | PVRL3 | 5366014798 | 686606001 | SNAP23 | 5836164157 | 639191647 |
| PVRL3 | 6295317014 | 780071609 | SH3BP4 | 5016521454 | 35729801 | SH3BP4 | 5759361267 | 394753858 |
| STEAP3 | 6276416779 | 52850408 | STEAP3 | 4899868011 | 473234716 | PVRL3 | 5714940707 | 735455653 |
| RASAL2 | 6018846194 | 540622251 | ITGB5 | 4856338819 | 344446616 | ACTBL2 | 5593639692 | 35834398 |
| ROBO1 | 5927528381 | 773979176 | UACA | 4774085681 | 536368122 | RBMX;RBMXI | 5555875142 | 62318085 |
| SH3BP4 | 59149278 | 401411459 | ACACB | 4693423589 | 550050498 | CD151 | 5505231222 | 2976925 |
| CD151 | 5906514486 | 311146505 | DBT | 4683039029 | 492270189 | NOTCH2 | 5502668381 | 555692566 |
| SLC30A1 | 5706652959 | 318988497 | TXNL1 | 4602086385 | 402943839 | VANGL1 | 549691232 | 552111723 |
| VANGL1 | 5705279986 | 468979087 | ABCC1 | 4568278631 | 47135967 | ROBO1 | 5355998357 | 771799129 |
| ZDHHC5 | 5678084691 | 69204001 | RPL36 | 4532576243 | 332656917 | PVRL2 | 5245352745 | 527179282 |
| HLA-A | 5624280294 | 500344629 | NOTCH2 | 4514105479 | 495769296 | RASAL2 | 5157117526 | 52175914 |
| VAMP5 | 5596427282 | 299471596 | CD151 | 447731336 | 246989908 | ZDHHC5 | 5124977112 | 659033452 |
| PVRL2 | 5567234993 | 566385929 | SLC30A1 | 4431339264 | 26899896 | SLC30A1 | 5124522527 | 304976538 |
| ITGB5 | 5478455544 | 341152107 | SLC4A7 | 4430386861 | 522273439 | BSG | 4978650411 | 509797313 |
| DAG1 | 5477238973 | 655922889 | ROBO1 | 4386838277 | 643847222 | STEAP3 | 494482549 | 413822999 |
| PPFIBP1 | 5459542592 | 359800913 | SNAP23 | 4263252576 | 51917233 | VAMP5 | 4869506836 | 266368029 |
| CDC42EP1 | 5449122111 | 699770827 | PVRL2 | 4061273257 | 472000166 | PHACTR4 | 4867570559 | 472090994 |
| PHACTR4 | 5328099251 | 49772584 | PPFIBP1 | 4024754206 | 274132417 | DCBLD2 | 4850252469 | 65168675 |
| SCRIB | 5322535515 | 368944121 | KIRREL | 3991343498 | 598634518 | CDC42EP1 | 4821839015 | 649255305 |
| DCBLD2 | 5291316032 | 690266022 | ZDHHC5 | 396969986 | 564281147 | HLA-A | 4804843903 | 455490629 |
| KIRREL | 5057974815 | 670088235 | ASPH | 3947157224 | 707748124 | PPFIBP1 | 4705691655 | 322358521 |
| SLC7A5 | 504317983 | 250244536 | VAMP5 | 3940592448 | 219466554 | SLC7A5 | 4668615341 | 233079653 |
| SHB | 5026381811 | 670600915 | SCRIB | 3904259046 | 288225256 | SNX33 | 4636424383 | 443198295 |
| SLC9A3R2 | 5014690399 | 494118229 | RPS5 | 3845104218 | 649860583 | CD99 | 4587390582 | 657668328 |
| UACA | 5005984306 | 556541351 | CDC42EP1 | 3833697001 | 596647085 | RELL1 | 4560972532 | 533755422 |
| RASA3 | 4968805949 | 481652441 | RASAL2 | 3831459999 | 377561352 | PKN2 | 4539492607 | 526089144 |
| FERMT2 | 4909849485 | 527675563 | ITGA5 | 3787483215 | 381036768 | FERMT2 | 4501648903 | 499969333 |
| RELL1 | 4881906192 | 545158629 | EFNB2 | 3780033429 | 54253062 | DAG1 | 4478713989 | 560787983 |
| ATP2B4 | 4844647408 | 540223164 | PAK4 | 3735708555 | 246243274 | SCRIB | 4458333015 | 323740318 |
| SLC3A2 | 4804778417 | 557583488 | CSPG4 | 3733531952 | 469059936 | KIRREL | 4383095741 | 600345225 |
| PARD3 | 4800222079 | 644568649 | CANX | 3732142448 | 421970622 | PAK4 | 4357777596 | 282362431 |
| PKN2 | 4790421486 | 534313117 | RELL1 | 3722721418 | 470402612 | EFNB2 | 4304113706 | 595167697 |
| EFNB2 | 4783253034 | 636450818 | RPL26;RPL26I | 3695710818 | 173786197 | PARD3 | 4301913897 | 640740892 |
| SLC4A7 | 4764970144 | 543715982 | RPL38 | 3630520821 | 244965908 | DIAPH3 | 4271723747 | 350142824 |
| RAB27B | 4751863798 | 542227609 | DCBLD2 | 3592331886 | 579302839 | GPR176 | 4268102328 | 541960565 |
| MARK2 | 4749516169 | 513585651 | SEMA7A | 3589494705 | 169476781 | ATP2B4 | 4250353813 | 502061996 |
| CSPG4 | 4714560827 | 524479427 | PTPRJ | 3496129036 | 236106679 | RPS26;RPS26 | 4249290466 | 218291568 |
| PAK4 | 4666082064 | 300197147 | YES1 | 3494282722 | 465635088 | SNX9 | 4245838483 | 557055442 |
| JAG1 | 4656602859 | 602551905 | EFR3A | 3463176409 | 424225664 | ITGA3 | 4219128927 | 742557738 |
| MARK3 | 4596480687 | 391585038 | RAC1;RAC3;R | 3425718307 | 399076441 | RGL2 | 4175549189 | 493344852 |
| VEPH1 | 4538318316 | 613177604 | BSG | 3420258204 | 401520577 | SLC4A7 | 4137465159 | 49867856 |
| ROCK1 | 4523753802 | 482414269 | FN1 | 3410116831 | 412636482 | RASA3 | 4128130595 | 422245661 |
| ITGA3 | 4451560338 | 784257184 | SHB | 3390035629 | 525173273 | SLC3A2 | 4073179245 | 507614063 |
| EFR3A | 4434311867 | 490258561 | GPR176 | 3389539401 | 474419575 | CDCA3 | 4066978455 | 502692204 |
| LLGL1 | 4421529134 | 759103738 | SLC7A5 | 3387625376 | 167760621 | PTPN14 | 4065708478 | 396216094 |
| GPR176 | 4404317538 | 546339962 | TCEB2 | 3367448171 | 194039632 | LLGL1 | 4018285116 | 742680018 |
| PCDH7 | 4390195211 | 82840752 | PARD3 | 3356177966 | 540671874 | SIRPA;SIRPB1 | 4013646444 | 524097547 |
| KIAA1522 | 4380814552 | 517496106 | SNTB1 | 3350037257 | 452257424 | TMEM2 | 3974522591 | 414444306 |
| CDCA3 | 4368008296 | 528998004 | PALM2 | 322900486 | 441838668 | PTPRJ | 3967389107 | 264075982 |
| PTPRJ | 4362202644 | 29087165 | FAM171B | 31888237 | 453564226 | ITGA7 | 396243 | 555170202 |
| YES1 | 4357068062 | 603220461 | DLG5 | 3185515086 | 326042444 | VEPH1 | 3950240771 | 601160782 |
| ANTXR2 | 4352134705 | 487730924 | PHACTR4 | 3164863269 | 347388392 | SHB | 3935052236 | 506585407 |
| ITGA6 | 4349503835 | 485778004 | SIRPA;SIRPB1 | 3138269424 | 425261298 | EFR3A | 3900314013 | 454990248 |
| USP6NL | 4340977987 | 613026422 | ITGA3 | 3112800598 | 695154799 | RPL38 | 389615345 | 253995705 |
| CDC42BPA | 4321214676 | 427432999 | SPECC1 | 3088081042 | 402438689 | CSPG4 | 3892767588 | 447286647 |
| SNX9 | 4319847425 | 573471655 | FERMT2 | 3060415586 | 394678916 | KIAA1522 | 3872991244 | 504013651 |
| MPZL1 | 4317663193 | 530675715 | RPL27 | 3058577855 | 19084972 | SNX18 | 3852434794 | 584652528 |
| PACSIN2 | 4260470708 | 604881793 | CDC42BPA | 3050934474 | 295416312 | MARK2 | 3816738129 | 447509025 |
| ITGA7 | 4256843249 | 610305328 | ATP2B4 | 3046378136 | 405990388 | SLC9A3R2 | 3801883698 | 414707767 |
| RGL2 | 4253754934 | 502213842 | RPS18 | 3040205638 | 26098869 | ITGA6 | 3786978404 | 446878039 |
| CASKIN2 | 4248947144 | 590277225 | SLC3A2 | 3017041524 | 426142045 | MPZL1 | 3764019648 | 492332591 |
| ITGB1 | 4240039508 | 867745274 | RPL35 | 3015224775 | 125914099 | FKRP | 3751907667 | 184573634 |
| PCDH10 | 4215988159 | 50890749 | DOCK10 | 3014827728 | 3471788 | ITGB1 | 3746035576 | 778921706 |
| DOCK10 | 4194951375 | 415380797 | JAG1 | 3013652166 | 46291922 | IL6ST | 3738710721 | 50654051 |
| CTNNA1 | 4161563873 | 429967916 | RPLP1 | 298711141 | 103005188 | YES1 | 3692814827 | 546308237 |
| CTNND1 | 4154057503 | 661150856 | NT5E | 2976449013 | 185136543 | CTNND1 | 3690487862 | 623458568 |
| SIRPA;SIRPB1 | 4149618785 | 515436951 | GNB2 | 2962380091 | 195680277 | UACA | 3689159711 | 46478484 |
| PTPN14 | 4140806516 | 403707406 | RPL34 | 2953519821 | 174519708 | NF2 | 3677895228 | 411020802 |
| RALGAPA1 | 4113625526 | 431472885 | MARK3 | 2941506068 | 254405285 | USP6NL | 3665531158 | 550703688 |
| CDC42EP4 | 4087483724 | 454331896 | HSPD1 | 2926898321 | 22582487 | CASK | 3659744898 | 418157997 |

| | | |
|---|---|---|
| IL6ST | 404711628 | 521516185 |
| BSG | 4037142436 | 377111891 |
| TENM3 | 4010353724 | 592767446 |
| RICTOR | 4009146055 | 555248756 |
| MXRA8 | 3981778145 | 543588576 |
| TXNL1 | 3978030523 | 370328736 |
| SLC39A6 | 3976314545 | 621031353 |
| SLC12A2 | 3964370728 | 638807863 |
| CD44 | 3960669835 | 627400529 |
| NF2 | 3957550049 | 428086739 |
| AHCYL1 | 395117569 | 245838023 |
| CPNE8 | 3921220462 | 530959756 |
| KCNMA1 | 392121315 | 541264731 |
| SLC7A11 | 3881628672 | 432867519 |
| SPECC1 | 3868702888 | 439077255 |
| DIAPH3 | 3857059797 | 314917012 |
| EVI2B | 3856755575 | 450674819 |
| PLSCR3 | 3848349253 | 63230738 |
| SNTB1 | 3841130575 | 425584648 |
| FMNL3 | 3828826586 | 414681211 |
| SLC39A10 | 3812507629 | 48259241 |
| HLA-C | 3801843961 | 346140788 |
| ADAM9 | 3766999563 | 462550326 |
| ANKRD50 | 3759481112 | 667283913 |
| SNX18 | 3693138758 | 578749195 |
| SNX33 | 3662002563 | 375015504 |
| PRR16 | 3635044734 | 509582724 |
| EHD2 | 3613792102 | 293489528 |
| FGD6 | 3589107831 | 488509783 |
| GAB1 | 3541888237 | 550541558 |
| TMEM2 | 3535220146 | 370144877 |
| ZDHHC20 | 3529731115 | 555213507 |
| ERBB2 | 3527312597 | 596484875 |
| C2CD2 | 3521382332 | 461085621 |
| EGFR | 3515865326 | 683838537 |
| SLITRK5 | 3492024422 | 364211634 |
| MTMR1 | 3490453402 | 425192676 |
| PALM2 | 3474478722 | 439444583 |
| FCHO2 | 3447653135 | 572305165 |
| VASN | 3433085442 | 51955394 |
| ATP2B1 | 3427261353 | 349656693 |
| GPRIN1 | 3425520897 | 246155743 |
| RAB23 | 3416476885 | 515428494 |
| GULP1 | 3411631266 | 263275668 |
| ROR1 | 3397293409 | 640269567 |
| SLC16A3 | 3374832153 | 547576635 |
| LPHN2 | 3365814845 | 359891621 |
| JUP | 3355384827 | 520635129 |
| MAP4K5 | 3339738528 | 57376006 |
| CTNNB1 | 3317666054 | 414221019 |
| WIPF2 | 3315406481 | 447632423 |
| WDR20 | 3291771571 | 515097499 |
| SLC20A1 | 3291093508 | 381385221 |
| PLXNA1 | 3286937078 | 427841945 |
| ESYT2 | 3280861219 | 301857673 |
| SLC39A14 | 3279722214 | 528834564 |
| FLOT1 | 3278645198 | 231148488 |
| ARHGAP32 | 3247563998 | 426442233 |
| SBF1 | 3226476351 | 56135348 |
| USP12 | 3218187968 | 457344024 |
| RASA2 | 3197110812 | 375107899 |
| KIAA1549L | 3196724256 | 51108799 |
| ABCC1 | 3183499018 | 36888386 |
| SLC1A5 | 3172509829 | 4264695 |
| KIDINS220 | 3155676842 | 369686881 |
| DSG2 | 3149492582 | 726687972 |
| CXADR | 3148575147 | 406594957 |
| CDH2 | 307377402 | 572573836 |
| YKT6 | 3060578028 | 541596705 |
| PEAK1 | 3047127406 | 233302615 |
| STIM2 | 3041267713 | 46183023 |
| RHBDF1 | 3038778623 | 433531228 |
| PLCB3 | 3034277598 | 451664301 |
| PTPN13 | 302687041 | 337600862 |
| CASK | 302634112 | 248706191 |
| SLC4A8 | 3021724065 | 363703351 |
| DBT | 3016879082 | 290002083 |
| EPB41L5 | 3000426292 | 623340233 |
| EPS8 | 2996011734 | 43570934 |

| | | |
|---|---|---|
| RPS17 | 2923841159 | 50821476 |
| RPS13 | 2878815015 | 16393358 |
| MPZL1 | 2876598994 | 415030168 |
| ANTXR2 | 2867431005 | 367311646 |
| SLC9A3R2 | 2855993271 | 376520204 |
| RPS19 | 2839215914 | 276562536 |
| CTNND1 | 2825543404 | 52618867 |
| EPB41 | 2819589615 | 624318546 |
| VEPH1 | 2815087318 | 50543061 |
| DST | 2812931697 | 218150214 |
| DSG2 | 2799290021 | 732022202 |
| ADAM9 | 2780865987 | 374229554 |
| ITGB1 | 2764720599 | 758008928 |
| DDOST | 2762450854 | 459383022 |
| FLOT2 | 2753797849 | 168187175 |
| GNB1 | 2736635526 | 157148782 |
| PCDH10 | 2733088175 | 416479799 |
| CD44 | 2732317607 | 49635671 |
| C2CD2 | 2728356997 | 389627716 |
| KCNMA1 | 2723492304 | 432712579 |
| CASK | 2651903788 | 32780438 |
| RPL36AL | 2650666237 | 097278347 |
| ILF3 | 2648517291 | 433256239 |
| EHD2 | 2630783399 | 223432979 |
| DAG1 | 2622069677 | 269927286 |
| FMNL3 | 261613814 | 366303385 |
| RPS25 | 2603988965 | 197302674 |
| SLC39A10 | 2585576375 | 381744014 |
| CDC42 | 2582015673 | 257871926 |
| FCHO2 | 2566893578 | 47841816 |
| NF2 | 2547934214 | 304912677 |
| JUP | 2519610723 | 430859189 |
| H2AFY | 2513070742 | 198322866 |
| RPS10;RPS10 | 2511832555 | 245786738 |
| EPHA2 | 2509480158 | 574776504 |
| SLC39A6 | 2507164637 | 25635949 |
| RPL32 | 2475898107 | 17004684 |
| AKAP12 | 2446949323 | 205767385 |
| PLSCR3 | 2436248779 | 499367211 |
| RRAS2 | 243268617 | 156746382 |
| HLA-A | 2428389867 | 109111206 |
| FLOT1 | 2427451452 | 171780908 |
| RPS26;RPS26 | 2416365941 | 075291481 |
| CACNA2D1 | 2415474256 | 137435078 |
| PTPN14 | 2410697619 | 256881113 |
| GNG12 | 241059653 | 138054888 |
| UTRN | 2409901619 | 210523445 |
| ACACA | 240687116 | 650672397 |
| SLC12A2 | 2386033376 | 457804873 |
| PKN2 | 2369648298 | 346094102 |
| PTRF | 2363715808 | 161439131 |
| SLC16A3 | 2347569148 | 446932978 |
| SNX9 | 2324026426 | 392693553 |
| FGD6 | 2319455147 | 367171347 |
| HIST1H1B | 2312712351 | 106106759 |
| MCCC2 | 2307245255 | 30138609 |
| LGALS1 | 2297730128 | 115433737 |
| CAV1 | 2296662331 | 102867151 |
| RPL30 | 2287799199 | 429941822 |
| KANK1 | 2284746488 | 153578993 |
| ITGA6 | 2266092936 | 289098802 |
| EGFR | 2263411204 | 553466927 |
| PCCA | 2254234632 | 629819835 |
| RPL18 | 2250788371 | 09523485 |
| RPLP2 | 2230167389 | 117278077 |
| KIDINS220 | 2214800199 | 31853604 |
| SNTB2 | 2208288511 | 201827131 |
| PLAUR | 2203823725 | 136168638 |
| HYOU1 | 2190600713 | 345394461 |
| MARK2 | 2182050069 | 229338414 |
| ATP2B1 | 2160401662 | 235872206 |
| LLGL1 | 2148841222 | 344495621 |
| ARHGAP32 | 2132658323 | 311100801 |
| MXRA8 | 2131309827 | 367240792 |
| PCCB | 2126790047 | 535972843 |
| NCEH1 | 2110885938 | 318301068 |
| HNRNPC | 2102642695 | 158735733 |
| ITGA7 | 2095074654 | 349456707 |
| MCCC1 | 209132576 | 754877914 |

| | | |
|---|---|---|
| TENM3 | 3652631124 | 575681686 |
| PACSIN2 | 3646870931 | 557640088 |
| EHD2 | 3624016126 | 298821528 |
| HLA-C | 3558598518 | 328371242 |
| PLSCR3 | 3544906616 | 582904613 |
| CTNNA1 | 350117747 | 391861434 |
| AHCYL1 | 3463287354 | 216370856 |
| ASPH | 3457563718 | 499342271 |
| JAG1 | 3452807109 | 515828956 |
| PALM2 | 3439772606 | 446923388 |
| SLC39A6 | 3433312098 | 630256042 |
| ADAM9 | 3407339732 | 434345115 |
| C2CD2 | 3393978437 | 452577652 |
| SLC39A10 | 3386978149 | 46031905 |
| SNTB1 | 3386117935 | 492872428 |
| EPS8 | 3384273529 | 470087556 |
| MARK3 | 3379856745 | 314184292 |
| CDC42EP4 | 3349086126 | 391674921 |
| RPS5 | 3340472539 | 390177741 |
| DOCK10 | 3334749858 | 385389431 |
| CASKIN2 | 333277003 | 543939858 |
| SPECC1 | 3322882016 | 425494167 |
| KCNMA1 | 3288410823 | 492669312 |
| GAB1 | 3278434436 | 527582916 |
| CTNNB1 | 3273874601 | 447116012 |
| S100A6 | 3254982313 | 083130662 |
| OCC1 | 3231505394 | 453507682 |
| ATP2B1 | 3226772944 | 330993148 |
| EVI2B | 3222738902 | 400084398 |
| CXADR | 3218651136 | 379031395 |
| CD44 | 3217728297 | 550630055 |
| ROCK1 | 3211436272 | 390663414 |
| FRMD6 | 3200148265 | 297971653 |
| ANTXR2 | 3195392609 | 405467751 |
| MXRA8 | 3193855604 | 482651092 |
| FGD6 | 3182449023 | 454794997 |
| PRR16 | 3182425181 | 489881623 |
| RICTOR | 3142087619 | 482100278 |
| PCDH7 | 3131856283 | 72329487 |
| KCNN4 | 3115003586 | 550464089 |
| ZDHHC20 | 3113851547 | 482716559 |
| SDC1 | 3109981855 | 236301328 |
| RASA2 | 3107797941 | 381231401 |
| MTMR1 | 3091632525 | 387564809 |
| SLC16A3 | 3063244502 | 503602391 |
| EGFR | 3044035594 | 642297511 |
| FMNL3 | 3018063545 | 391143002 |
| ATP1B3 | 3017755191 | 395984706 |
| CDC42BPA | 3012154897 | 335293971 |
| FCHO2 | 2981986046 | 526897707 |
| SLC20A2 | 295830663 | 481663954 |
| SLC12A2 | 2943490346 | 549986715 |
| CDC42 | 2937186877 | 269802198 |
| MAP4K3 | 2932632128 | 501614588 |
| WIPF2 | 2930672646 | 403872588 |
| ECE1 | 2908926328 | 410363873 |
| SLC7A11 | 290855217 | 370849385 |
| MAP4K5 | 2900500933 | 524114525 |
| CDC42EP3 | 2894538244 | 56742247 |
| AMIGO2 | 2894410769 | 541501469 |
| ROR1 | 2891388257 | 619769302 |
| ABI2 | 2880473455 | 19395573 |
| PCDH10 | 2879110336 | 430463437 |
| WASL | 2862318357 | 214455833 |
| LZTS2 | 2853641192 | 518944341 |
| SLC1A5 | 2810548464 | 390345679 |
| ZDHHC8 | 2791353861 | 421805706 |
| ARF6 | 2787260373 | 164939687 |
| USP12 | 27647597 | 408505511 |
| MTMR10 | 2751226107 | 388699049 |
| WDR20 | 2730328878 | 499223145 |
| MAPKAP1 | 272151343 | 43349226 |
| FLOT1 | 2709981283 | 193038493 |
| SLC16A1 | 2702795982 | 161683025 |
| ARHGAP22 | 2652642886 | 416184138 |
| MT-CO2 | 2641728719 | 280561135 |
| JUP | 2630900065 | 43201256 |
| RAB23 | 2630801837 | 443739004 |
| SBF1 | 2613571485 | 500488726 |

| | | |
|---|---|---|
| ZDHHC8 | 2993091583 | 428200078 |
| FRMD6 | 2992075602 | 286998608 |
| CD99 | 2976977984 | 152271218 |
| DLG1 | 2964256287 | 433736077 |
| MET | 296031634 | 467934532 |
| UTRN | 2943888028 | 245631737 |
| FAT4 | 2928129832 | 369313767 |
| DLG5 | 2921272278 | 291295956 |
| VAMP3 | 2914047241 | 502132669 |
| PPFIA1 | 2900395075 | 322263532 |
| ARHGAP22 | 2899224917 | 472985687 |
| CLMP | 2893290202 | 500013229 |
| AKAP12 | 2884889921 | 229418711 |
| WASL | 2881099383 | 21918913 |
| MINK1 | 2862959544 | 433532137 |
| BAIAP2 | 2831197739 | 534321935 |
| DOS | 2809056282 | 605756987 |
| EPHA2 | 2805438995 | 59569163 |
| SLC20A2 | 2793762207 | 487055133 |
| PAG1 | 278295517 | 40598188 |
| SNTB2 | 2769619306 | 252489214 |
| VAT1 | 2768820763 | 435763455 |
| ITGA1 | 2724829674 | 651646024 |
| RPL26;RPL26I | 2724649747 | 126021622 |
| CDC42EP3 | 2696837107 | 57602262 |
| PHACTR2 | 268176047 | 466176572 |
| MAGI1 | 2676602046 | 335356542 |
| ARHGAP21 | 2665767352 | 180307647 |
| LZTS2 | 2655083656 | 421182249 |
| SLC16A1 | 2647906303 | 155878536 |
| CDC42BPB | 2641875267 | 300053317 |
| NF1 | 2639081001 | 259552906 |
| KIAA0754 | 2605723699 | 30345161 |
| SLC9A3R1 | 2599212011 | 200943603 |
| ARL13B | 258856678 | 617045506 |
| KCNN4 | 2578001022 | 508056599 |
| CCDC88A | 2563981056 | 17193303 |
| ERBB2IP | 2558638573 | 554863386 |
| ABI2 | 255259641 | 167629348 |
| PLEKHA5 | 2538347562 | 440359777 |
| CLDND1 | 2516362826 | 131650227 |
| EPB41L1 | 2468669891 | 286079844 |
| CEP89 | 2447237015 | 442951569 |
| FAM129B | 2402788798 | 490902206 |
| FLOT2 | 2389297803 | 119403849 |
| TBC1D8 | 2380848249 | 182668835 |
| TRIOBP | 2378250122 | 369076631 |
| PTRF | 2372019132 | 160868957 |
| BASP1 | 2351034164 | 443350448 |
| DOCK9 | 2346939087 | 373502338 |
| C1orf21 | 2341188749 | 331863285 |
| C2CD5 | 233877182 | 352563123 |
| PLCB1 | 2338770231 | 439934368 |
| CAV1 | 2316704114 | 103878251 |
| PSD3 | 2311279297 | 316912464 |
| DEPDC1B | 2307048162 | 451001201 |
| MTMR10 | 2305902163 | 238512757 |
| RAB11B;RAB1 | 2266668002 | 173968855 |
| NOTCH1 | 2260386467 | 173125839 |
| TBC1D10B | 2253235817 | 264044241 |
| BCAR3 | 2244311651 | 177670445 |
| TNFRSF11A | 2242758115 | 291166469 |
| OCC1 | 2227365812 | 145434746 |
| RAI14 | 2221817334 | 458614838 |
| EPB41 | 2197956085 | 545964668 |
| CYFIP2 | 2195943832 | 136513509 |
| TULP3 | 2194774628 | 222030395 |
| PI4KA | 2194368998 | 338241509 |
| EPB41L2 | 218880113 | 55884371 |
| MAP4K3 | 2184784253 | 142078879 |
| DLC1 | 2173642159 | 251585293 |
| ROCK2 | 2168944359 | 202188417 |
| MARCKS | 2162693659 | 541757269 |
| NUMB | 2155855497 | 691509174 |
| EPHA4 | 215104421 | 276095599 |
| EHD1 | 2126116435 | 427233221 |
| RAB34 | 2104054769 | 238850387 |
| FIBP | 2084970792 | 292311627 |
| ANKS1A | 2054703712 | 166462546 |

| | | |
|---|---|---|
| ROCK1 | 2083170891 | 262937843 |
| ERBB2IP | 2082267443 | 519970953 |
| SDC1 | 2076841672 | 081891535 |
| SLC1A5 | 2075257937 | 307185403 |
| TRIOBP | 2074475606 | 395506131 |
| DLG1 | 2049403509 | 330741452 |
| GNAI2 | 2040617943 | 188590728 |
| MYO1B | 2034568151 | 262750693 |
| RPL10A | 2019964854 | 459569313 |
| SLC4A8 | 2013962428 | 183611919 |
| CYB5R3 | 2005530039 | 10039186 |
| ALCAM | 200410525 | 142627542 |
| HIST1H2AJ;HI | 1996999423 | 238632139 |
| MINK1 | 1996735891 | 309895131 |
| RPL6 | 199150912 | 531457709 |
| IQGAP1 | 1975510915 | 262439166 |
| RASA3 | 1968233744 | 178507888 |
| PEAK1 | 1965897878 | 147347048 |
| SLITRK5 | 1963186582 | 230065794 |
| CD59 | 1953125318 | 186516104 |
| RPN1 | 1952860196 | 151608802 |
| APBB2 | 1949297905 | 165469527 |
| PHB | 1943523725 | 112303034 |
| EPB41L2 | 1931606611 | 524939346 |
| RICTOR | 1890750567 | 267755157 |
| ATP1A1 | 1886513392 | 132800405 |
| SBF1 | 188606294 | 327101414 |
| ACSL4 | 1869854291 | 141797039 |
| DLST | 1859298388 | 322033527 |
| RAB34 | 1842550278 | 208691092 |
| IPO5 | 1809966087 | 150047325 |
| CXADR | 1803896904 | 282121268 |
| HSP90B1 | 1793602308 | 169621206 |
| RAB11B;RAB1 | 1784909884 | 132550862 |
| FAM171A2 | 1772495588 | 164928904 |
| RPL7A | 1756951014 | 228494926 |
| ESYT2 | 1754687945 | 164978952 |
| FRMD6 | 1744161606 | 167923601 |
| RPLP0;RPLP0I | 173088328 | 364890391 |
| RPL8 | 1721759796 | 460227332 |
| PTPN13 | 1718651136 | 224579885 |
| RPL12 | 1718208631 | 530811303 |
| P4HA1 | 1713344256 | 164340775 |
| RAI14 | 1673179944 | 72939795 |
| RPL35A | 1652516365 | 405433035 |
| PC | 1638703028 | 416513193 |
| VAMP3 | 1584064484 | 341409658 |

| | | |
|---|---|---|
| MET | 2611097972 | 440420996 |
| DLG1 | 2605836233 | 389408093 |
| BCAR3 | 2596420288 | 222241297 |
| MAGI1 | 2590811412 | 323368218 |
| RPS18 | 2576444626 | 204826337 |
| DHX30 | 2558403651 | 474738595 |
| DLC1 | 2549741427 | 303228884 |
| TNS1 | 2549218814 | 278445297 |
| ESYT2 | 2546870867 | 240315859 |
| CPNE8 | 2546075503 | 395384796 |
| KIAA1549L | 2543934186 | 482793207 |
| TENM4 | 252050813 | 243657109 |
| PHACTR2 | 2519277891 | 512135538 |
| DSG2 | 2489291827 | 611596258 |
| EPB41L5 | 2487856229 | 567622393 |
| RAB27B | 247818025 | 155514026 |
| PEAK1 | 2476418177 | 191706736 |
| LPHN2 | 2475931168 | 284379528 |
| CNP | 2467754046 | 272640153 |
| VASN | 2464906375 | 435195709 |
| VAT1 | 2454834938 | 419442053 |
| VAMP3 | 2434059143 | 461690004 |
| RAP1B;RAP1A | 2432903608 | 121843316 |
| BAIAP2 | 2431918462 | 484157955 |
| FAM171B | 2431312561 | 364493459 |
| YKT6 | 2429345449 | 410452 |
| GULP1 | 2422508876 | 194763188 |
| ALCAM | 2415833155 | 313882153 |
| PLCB3 | 2407239914 | 394103512 |
| RAB27A | 2399236997 | 195415464 |
| C2CD5 | 2397780736 | 362419562 |
| HSD17B12 | 2390179952 | 189085825 |
| CANX | 2380681038 | 164432562 |
| ROCK2 | 2377708117 | 301033447 |
| EPHA2 | 2345383962 | 526800859 |
| PPFIA1 | 2343330383 | 275017561 |
| ANKRD50 | 2299443245 | 367587972 |
| RAB10 | 2298195839 | 150753517 |
| VDAC3 | 2283137639 | 104794242 |
| RAB34 | 2279689789 | 254962677 |
| SLC25A4 | 2263328234 | 169150065 |
| PDGFRB | 2258238475 | 384233076 |
| RPS13 | 2255426089 | 112637514 |
| RPL36 | 2249342601 | 120147232 |
| DHCR24 | 2246846835 | 342941768 |
| ATP1A1 | 2239752452 | 161792826 |
| RAB11B;RAB1 | 2235394478 | 162804872 |
| TXNIP | 2224891345 | 287525903 |
| RPN1 | 2221980731 | 17000286 |
| ERBB2IP | 2219847679 | 545655477 |
| PTRF | 2207962354 | 149597412 |
| TMEM165 | 2200647036 | 148685152 |
| DOCK5 | 2187198321 | 351059985 |
| DOCK9 | 2184613546 | 361046127 |
| BASP1 | 2179946899 | 416716807 |
| ITSN2 | 2177964846 | 230390079 |
| TLDC1 | 2170835813 | 271346916 |
| YWHAG | 2169887861 | 144510699 |
| ARHGAP32 | 2164774895 | 359873068 |
| SAR1A | 2160915693 | 127357026 |
| FIBP | 2158139229 | 292356108 |
| LNPEP | 2153412501 | 339426491 |
| DLG5 | 2135985057 | 230625623 |
| TM9SF3 | 212778314 | 135729649 |
| EHD4 | 2104756991 | 531241237 |
| SLC4A8 | 209705925 | 269249158 |
| RPS25 | 2074393272 | 140292741 |
| TBC1D10B | 2070303281 | 256679134 |
| CYFIP2 | 2059843063 | 129132028 |
| SNTB2 | 2057745934 | 187033879 |
| UTRN | 205682977 | 179408133 |
| SLC20A1 | 2051305453 | 13531979 |
| RPN2 | 2044282595 | 15674745 |
| PTPN13 | 2043244998 | 251954612 |
| SLC9A3R1 | 2030046463 | 150859688 |
| ABI1 | 2026866595 | 382538569 |
| CEP89 | 2020237605 | 461959732 |
| APBB2 | 2013462385 | 171449161 |
| HSPD1 | 1984737396 | 139757783 |

| | | |
|---|---|---|
| ZC3HAV1 | 2012102763 | 324246195 |
| KIAA1217 | 2001148542 | 158800266 |
| MCCC2 | 1988578161 | 228279284 |
| RRAS2 | 1963589668 | 120046666 |
| ADD3 | 1948035876 | 432117773 |
| MAP4K4 | 1944439252 | 349066534 |
| ATP1A1 | 1927450816 | 135088259 |
| DOCK5 | 1920156161 | 315059301 |
| APBB2 | 1910768827 | 160827165 |
| SPRED2 | 1890294393 | 275012997 |
| SLC29A1 | 186755085 | 121908131 |
| FAM171B | 1863147736 | 152192756 |
| ABI1 | 185888958 | 31475451 |
| KIAA1549 | 1846309026 | 148985068 |
| MYOF | 1818333626 | 352377067 |
| RPS27A;UBA5 | 1816491763 | 458909334 |
| ADD1 | 1813187281 | 451009529 |
| RASSF8 | 1807260195 | 117539252 |
| ARHGAP39 | 1805556615 | 547237346 |
| FNBP1 | 1800481478 | 162694073 |
| SNAP29 | 1793507576 | 546001685 |
| ASAP1 | 1749130567 | 453233419 |
| TP53BP2 | 1741802533 | 136753485 |
| STXBP1 | 1738177617 | 307918586 |
| C2CD2L | 1737789154 | 256040004 |
| SRGAP1 | 1728999774 | 143107398 |
| AMIGO2 | 1696874936 | 139543295 |
| LIN7C | 1681933403 | 118329111 |
| SIPA1L3 | 167793115 | 136365618 |
| EPS15 | 1677246094 | 198877794 |
| CDK5 | 1667403221 | 316703459 |
| MPP5 | 1648251851 | 265471237 |
| CYFIP1 | 164294974 | 398910113 |
| CDC42 | 1637461027 | 122399792 |
| COBLL1 | 1622813861 | 121814406 |
| WASF2 | 1620409012 | 397804112 |
| DLGAP4 | 1610899289 | 145124119 |
| NCKAP1 | 1593482335 | 503263938 |
| DAB2 | 1561963081 | 538019714 |

| | | |
|---|---|---|
| ERBB2 | 1972148895 | 489013488 |
| XPO1 | 1952496847 | 197246272 |
| DNAJA1 | 1933801651 | 277072799 |
| KIAA0754 | 1906539917 | 256808056 |
| PLEKHA5 | 1898013115 | 383628059 |
| EPB41L2 | 1877187411 | 507117559 |
| MARCKS | 1864081701 | 546778909 |
| ANKS1A | 185272185 | 150593293 |
| CDH2 | 1844578743 | 155005845 |
| SLC12A4 | 1839426676 | 27953561 |
| FRS2 | 1833919843 | 324714615 |
| NCEH1 | 1831219037 | 161181971 |
| FAM129B | 1816116333 | 435887958 |
| AKAP12 | 1812652906 | 162515882 |
| ASAP1 | 1808398883 | 462162331 |
| TRAM1 | 1799631755 | 180797879 |
| UQCRC2 | 1780291557 | 20091002 |
| DNAJA2 | 1773986816 | 369429445 |
| DLGAP4 | 175554053 | 175190804 |
| STIM2 | 1755379041 | 178047229 |
| ARL3 | 1738911311 | 217146659 |
| EHD1 | 173078537 | 375147162 |
| PI4KA | 1727696101 | 290933934 |
| CYFIP1 | 1672404289 | 51192116 |
| ARHGAP39 | 1656159083 | 575125871 |
| WASF2 | 1651583989 | 508196167 |
| NUMB | 1650914828 | 63456897 |
| NCKAP1 | 1593406677 | 61754125 |
